# Predictive coding models for pain perception

**DOI:** 10.1101/843284

**Authors:** Yuru Song, Mingchen Yao, Helen Kemprecos, Áine Byrne, Zhengdong Xiao, Qiaosheng Zhang, Amrita Singh, Jing Wang, Zhe S. Chen

**Affiliations:** Department of Psychiatry, New York University School of Medicine, New York, United States; Department of Biology, University of California, San Diego, United States; Kuang Yaming Honors School, Nanjing University, Nanjing, China; Department of Biochemistry, New York University, New York, United States; Center for Neural Science, New York University, New York, United States; Department of Anesthesiology, Pain and Operative Medicine, New York University School of Medicine, New York, United States; Department of Neuroscience and Physiology, New York University School of Medicine, New York, United States; Neuroscience Institute, New York University School of Medicine, New York, United States

**Author notes:** **These authors contributed equally** (Yuru Song, Mingchen Yao). **Corresponding, Correspondence: Z. S. Chen, Department of Psychiatry, New York University School, of Medicine, New York, NY 10016, USA, **.

**Keywords:** Predictive coding, pain perception, somatosensory cortex, anterior cingulate cortex, mean field model, placebo, chronic pain

## Abstract

Pain is a complex, multidimensional experience that involves dynamic interactions between sensory-discriminative and affective-emotional processes. Pain experiences have a high degree of variability depending on their context and prior anticipation. Viewing pain perception as a perceptual inference problem, we propose a predictive coding paradigm to characterize evoked and non-evoked pain. We record the local field potentials (LFPs) from the primary somatosensory cortex (S1) and the anterior cingulate cortex (ACC) of freely behaving rats—two regions known to encode the sensory-discriminative and affective-emotional aspects of pain, respectively. We further use predictive coding to investigate the temporal coordination of oscillatory activity between the S1 and ACC. Specifically, we develop a phenomenological predictive coding model to describe the macroscopic dynamics of bottom-up and top-down activity. Supported by recent experimental data, we also develop a biophysical neural mass model to describe the mesoscopic neural dynamics in the S1 and ACC populations, in both naive and chronic pain-treated animals. Our proposed predictive coding models not only replicate important experimental findings, but also provide new prediction about the impact of the model parameters on the physiological or behavioral read-out—thereby yielding mechanistic insight into the uncertainty of expectation, placebo or nocebo effect, and chronic pain.

## 1 INTRODUCTION

Pain is a fundamental experience that is subjective and multidimensional. Pain processing involves sensory, affective, and cognitive processing across distributed neural circuits (Bushnell et al., 1999; Bushnell et al., 2013; Iannetti & Mouraux, 2010; Legrain et al., 2011; Vierck et al., 2013). However, a complete understanding of pain perception and cortical pain processing has remained elusive. Given the same nociceptive stimuli, the context matters for pain percept. In the literature, human neuroimaging studies have shown that among many brain regions, the primary somatosensory cortex (S1) and the anterior cingulate cortex (ACC) are two important cortical areas involved in high-level pain processing. The S1 represents the sensory-discriminative component of pain, whereas the ACC represents the affective-motivational component of pain (Johansen et al., 2001; Bushnell et al., 2013). In addition, ACC neuronal activities have been shown to correlate with noxious stimulus intensities, and chronic pain can alter acute pain intensity representation in the ACC to increase the aversive response to noxious stimuli at anatomically unrelated sites (Zhang et al., 2017). In addition to the bottom-up input, top-down attention, expectation, or contextual factors can bias cortical pain processing or modulate the strength or salience of pain signals (Wiech, 2016). Descending modulation can attenuate the incoming nociceptive signal and further skew the subjective pain perception despite the high-intensity noxious stimulus input.

Evoked pain is triggered by noxious sensory stimuli, whereas spontaneous pain (also known as non-evoked pain or non-evoked nociception) is not. Spontaneous pain can be induced by repeated noxious stimulations in naive animals, or induced by chronic pain conditions (Bennett, 2012). Pain perception has been conceptualized as perceptual inference (Wiech, 2016; Geuter et al., 2017; Tabor et al., 2017), and predictive coding may provide a theoretical model for characterizing such inference (Arnal & Giraud, 2012; Ploner et al., 2017). Specifically, pain perception can be studied as an inferential process in which prior information is used to generate expectations about future perception and to interpret the sensory input.

Predictive coding paradigms describe the inversion of a generative model of the percept and constantly adapt the hypothesis of sensory perception (Huang & Rao, 2011). A predictive model characterizes the uncertainty of sensory inputs in time and space (Aitchison & Lengyel, 2017). Predictive coding relies on correcting errors resulting from comparisons between internal predictions and actual observations. Such paradigms have provided important insights into perceptual inference, sensory processing, motor control, multi sensory integration and pain (Rao and Ballard, 1999; Shipp et al., 2013; Talsma, 2015; Sedley et al., 2016; Morrison et al., 2013; Hoskin et al., 2019). Predictive coding has been suggested as a universal computational principle in the neocortex (Bastos et al., 2012; Friston & Kiebel, 2009), and this framework may accommodate various data modalities and multiple spatiotemporal scales (Friston et al., 2015).

The experience of pain is often associated with brain rhythms or neuronal oscillations at different frequencies (Ploner et al., 2017; Peng et al., 2018). For multisite recordings, it is important to investigate the inter-regional local field potential (LFP) oscillatory coordination (Eto et al., 2011), as interareal oscillatory synchronization plays an important role in top-down neocortical processing (Bressler & Richtler, 2015; Bastos et al., 2020). One important theoretical implication of predictive coding is spectral asymmetries between the bottom-up and top-down representations (Bastos et al., 2012). The spectral asymmetry can be also explained by the functional asymmetry: prediction errors (PEs) express higher frequencies than the predictions that accumulate them, whereas the conversion of PEs into predictions entails a loss of high frequencies. Since the common characteristic frequencies in predictive coding range between the beta and gamma frequency bands, one working hypothesis is that the bottom-up PEs are represented at the gamma band and top-down prediction predictions are represented at the beta band.

In a series of rodent pain experiments, we collected various *in vivo* neurophysiological recordings from single or two brain regions in freely behaving rats (Zhang et al., 2017; Urien et al., 2018; Dale et al., 2018; Xiao et al., 2019). These data have established the foundation for improved understanding of pain perception and provided empirical evidence for computational modeling. In this paper, we present a predictive coding framework to model the temporal coordination of interareal oscillatory activity between the rat S1 and ACC during evoked and non-evoked nociception episodes. Specifically, we develop two different computational models to reproduce some previously observed differences between gamma and beta responses, before and after pain. The first model bears a form of the state space model based upon predictive coding, which can predict experimentally observed LFP responses at the gamma and beta bands in the S1 and ACC areas, respectively. The second model is derived from the mean field model, which is a biologically plausible neural mass model parameterized in terms of connection strengths between distinct neuronal subpopulations. The neural mass model can predict the S1 and ACC population neuronal activity in various pain conditions, for both naive and chronic pain-treated animals.

Our key hypothesis is that we can reproduce empirical findings by manipulating the gain parameter of the predictive coding model. Furthermore, the same phenomena can be reproduced by varying the synaptic efficacy in the neural mass model. In other words, we hypothesize that synaptic efficacy within the cortical pain network is a sufficient explanation for responses induced by pain, and variations in pain conditions correspond to variations in the model parameters described in the predictive coding paradigm.

In the result section, we first summarize important experimental findings that are extracted from previous published data (Xiao et al., 2019; Singh et al., 2020), which provide the biological support and motivation for our computational modeling work. We then describe our phenomenological model and mean-field model and their simulation results for both evoked pain and non-evoked nociception. Specifically, we will adapt the mean-field model to characterize pain aversive behaviors in chronic pain. We will make data interpretation and prediction related to the experimental results. To the best of our knowledge, this is the first systematic modeling investigation towards understanding pain perception. Together, our two computational models provide new insights into the uncertainty of expectation, placebo or nocebo effect, and chronic pain.

## 2 METHODS

### 2.1 Experimental Protocol and Recordings

All experimental studies were performed in accordance with the National Institutes of Health (NIH) *Guide for the Care and Use of Laboratory Animals* to ensure minimal animal use and discomfort, and were approved by the New York University School of Medicine (NYUSOM) Institutional Animal Care and Use Committee (IACUC).

Male adult Sprague-Dale rats (250-300 g, Taconic Farms, Albany, NY) were used in our current study and kept at the new Science Building at NYUSOM, with controlled humidity, temperature and 12-h (6:30 a.m.-6:30 p.m.) light-dark cycle. Food and water were available ad libitum. Animals were given on average 10 days to adjust to the new environment before the initiation of experiments.

Thermal pain stimuli were used for freely exploring rats in a plastic chamber of size 38 × 20 × 25 cm^3^ on top of a mesh table. A blue (473 nm diode-pumped solid-state) laser with varying laser intensities was consistently delivered to the rat’s right hindpaw. The laser stimulation (with intensity ranging 100-250 mW) was delivered in repeated trials (25-40) during 30-45 min. Two video cameras (120 frame per second) were used to continuously monitor the rat’s behavior during the course of experiment. Five naive rats and two chronic pain-treated rats were used in the current study. Details are referred to previous publications (Zhang et al., 2017; Dale et al., 2018).

To produce chronic inflammatory pain, 0.075 ml of *Complete Freund’s adjuvant* (CFA) (my-cobacterium tuberculosis, Sigma-Aldrich) was suspended in an oil-saline (1:1) emulsion, and injected subcutaneously into the plantar aspect of the hindpaw opposite to the paw that was stimulated by a laser; namely, only a unilateral inflammation was induced. In CFA rats, laser stimulations were delivered to the opposite paw of the injured foot. The ACC and S1 electrodes were implanted on the contralateral side of the stimulated foot.

Repeated noxious laser stimulations to the rat hindpaw could induce spontaneous pain behaviors. During the inter-trial intervals, we examined the rat’s behavior to identify non-evoked nociception episodes (such as twitch, lifting/flicking, paw withdrawal and paw licking) (Xiao et al., 2019).

We used silicon probes (Buzsaki32, NeuroNexus) with a 3D printed drive to record multichannel (up to 64 channels) neural activities from the rat ACC and S1 areas simultaneously, on the contralateral side of the paw that received noxious stimulation. For surgery, rats were anesthetized with isoflurane (1.5%-2%). The skull was exposed and a 3 mm-diameter hole was drilled above the target region. The coordinates for the ACC and S1 implants were: ACC: AP 2.7, ML 1.4-2.0, DV 2.0, with an angle of 20° toward the middle line; S1: AP −1.5, ML 2.5-3.2, DV 1.5. The drive was secured to the skull screws with dental cement. We used a Plexon (Dallas, TX) data acquisition system to record *in vivo* extracellular neural signals at a sampling rate of 40 kHz. The signals were first band-pass filtered (0.3 Hz-7.5 kHz), and LFPs were obtained upon subsequent band-pass filtering (1-100 Hz).

### 2.2 Data Analysis

#### Time-frequency analyses

Based on the simultaneously recorded multichannel LFP signals from the S1 and ACC, we applied the principal component analysis (PCA) and extracted the dominant principal component (PC) for the S1 and ACC, respectively. We then computed the spectrogram of the PC for each region. Multitapered spectral analyses for LFP spectrogram were performed using the Chronux toolbox (chronux.org). Specifically, we chose a half-bandwidth parameter *W* such that the windowing functions were maximally concentrated within [−*W, W*]. We chose *W* > 1*/T* (where *T* denotes the duration) such that the Slepian taper functions were well concentrated in frequency and had bias reducing characteristics. In terms of Chronux function setup, we used the tapers setup [*TW, N*], where *TW* is the time-bandwidth product, and *N* = 2 × *TW* − 1 is the number of tapers. Since the taper functions are mutually orthogonal, they give independent spectral estimates. In all time-frequency analyses, we used a moving window length of 500 ms and a step size of 1 ms. We used *TW* = 5. From the spectrogram, we computed the Z-scored spectrogram, where the baseline was defined as the 5-s period before the stimulus presentation.

#### Pain-responsive neurons

To identify pain-responsive neurons, we used a previously established criterion (Dale et al., 2018). Specifically, we computed the Z-scored firing rate related to the baseline (3-5 s before the stimulus onset). A neuron was called a positive pain-responding neuron if the following two criteria were satisfied: (i) the absolute value of the Z-scored firing rate of least one time bin (i.e., 50 ms) after stimulation must be greater than 2.5, and (ii) if the first criterion is met, at least the next two bins (i.e., 100 ms) must be greater than 1.65. These criteria must be fulfilled within 3 s after the stimulus onset.

#### Z-scored LFP power analysis

From the recorded multichannel LFPs of the S1 and ACC, we computed the Z-scored spectrogram for pain episodes (time 0 represents the laser onset in evoked pain, and the withdrawal onset in non-evoked nociception). During evoked pain, we usually observed event-related potentials (ERPs) in both S1 and ACC areas. Our prior report has indicated that the ERP latency was sooner (~200-300 ms) in the S1 than in the ACC during evoked pain episodes (Xiao et al., 2019). In contrast, during non-evoked nociception episodes, ERPs occurred in either the S1, or ACC, or both areas, with a high degree of variability in latency.

For non-evoked nociception episodes, we investigated whether the LFP power in the ACC and S1 at the beta and/or gamma bands change in a temporally coordinated manner. We computed the 10-s LFP spectrograms centered around the non-evoked nociception behavior onsets (pre-event: [−5, 0] s, post-event: [0,5] s). To highlight the event-related synchronization/desynchronization (ERS/ERD) phenomenon, we computed the Z-scored pre-gamma power related to the post-event period, and computed the Z-scored post-beta power related to the pre-event period.

Unless stated otherwise, all statistical tests were nonparametric tests without the normality assumption.

### 2.3 A Framework of Predictive Coding for Pain

#### Background

Predictive coding is aimed at minimizing the PE and further using it to update the prediction. The schematic diagram of predictive coding is shown in Fig. 1. To explain the predictive coding idea, we first introduce some basic notations. Specifically, let the latent variable *z* denote the subjective pain percept, let *x* denote the stimulus input. We also assume that *u* and *v* are two response variables, which represent the proxy for the observed gamma activity from the S1 and the beta activity from the ACC, respectively.

**Fig. 1:**
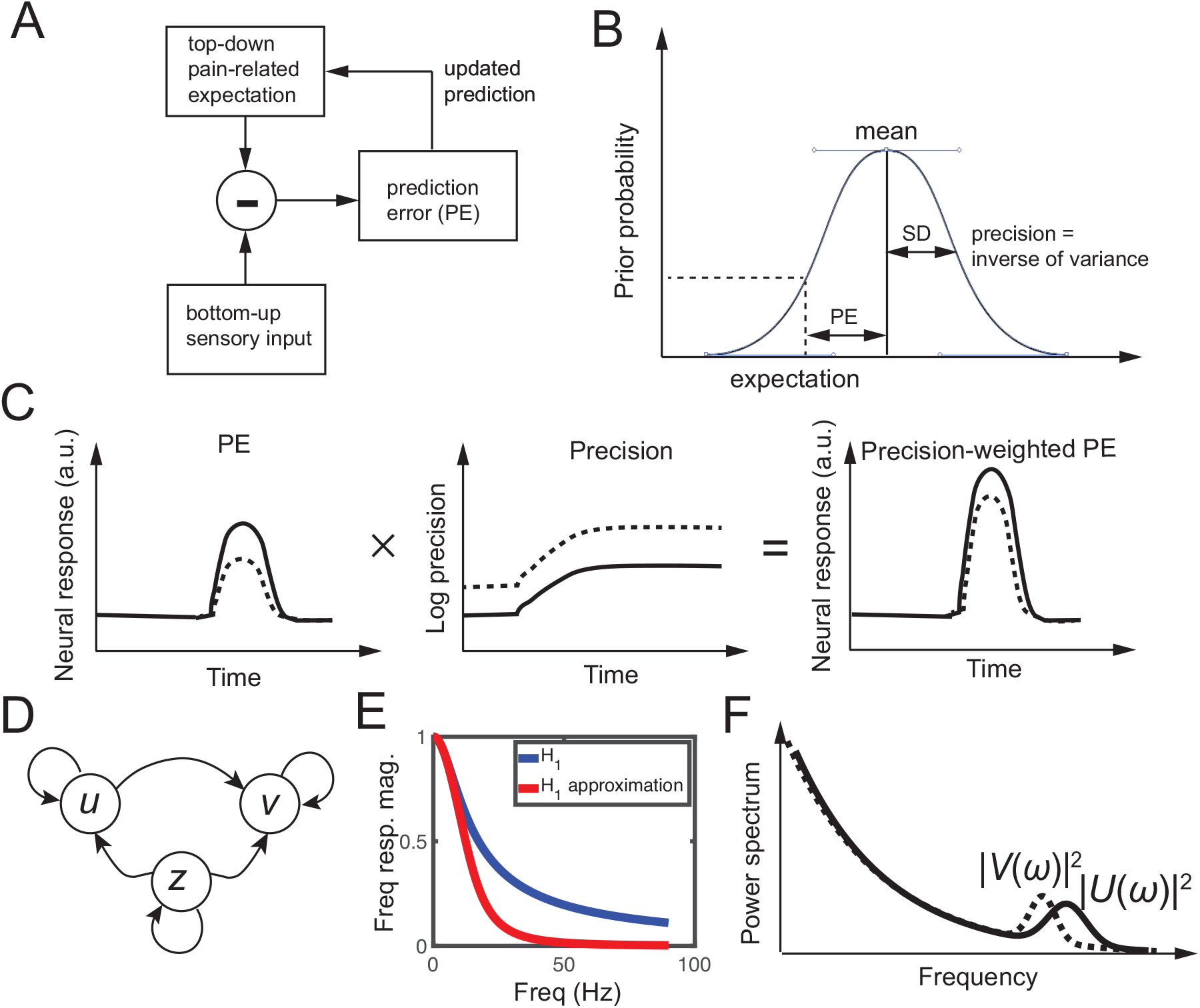
Predictive coding. (*A*) Schematic diagram of predictive coding for pain perception. (*B*) Graphical illustration of prediction error (PE), prior expectation, and prediction in perceptual inference. Due to the uncertainty of the top-down expectation, PE is assumed to be Gaussian distributed. Mean and standard deviation (SD) characterize the uncertainty of a Gaussian random variable. (*C*) Schematic illustration of neural response (a.u.) representing a gain-weighted PE that changes in time, where the gain is the precision statistic. (*D*) Graphical model showing statistical dependencies between the observed variables {*u, v*} and the latent variable {*z*} in the predictive coding model. Here, *z* denotes the pain expectation, and *u* and *v* denote the observed neural responses at two brain areas. (*E*) The magnitude of frequency response *H*_1_(*ω*) and its approximation, which can be viewed as a low-pass filter. (*F*) Schematic illustration of power spectra |*U* (*ω*)|^2^ (solid line) and |*V* (*ω*)|^2^ (dashed line), where *U* (*ω*) and *V* (*ω*) are the Fourier transforms of *u*(*t*) and *v*(*t*) in the predictive coding model, respectively.

In brief, predictive coding is used to dynamically update posterior expectations of pain (*z*) based upon PE. The underlying PEs and posterior expectations are then used to generate observable induced neural responses (*u* and *v*). To account for axonal conduction delays, we used stochastic delay differential equations for the predictive coding scheme.

#### Mathematical equations

First, we define the PE as the difference between the bottom-up finite-duration sensory input *x* and top-down pain-induced expectation *z* (Fig. 1*A*):

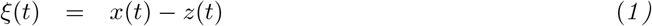

Predictive coding uses the signed PE to update the expectation after a certain time delay. Specifically, we assume that the dynamics of pain percept *z* follow a stochastic differential equation as follows

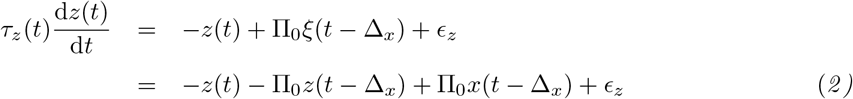

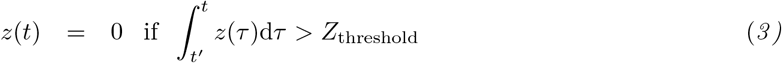

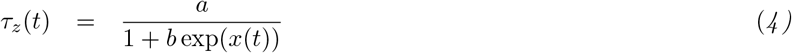

where Δ_*x*_ denotes a time delay parameter starting from the stimulus onset (Table 1), and *ϵ_z_* denotes the additive Gaussian noise. *Equation 2* is a linear delay-differential equation that characterizes the expectation update dynamics based on the PE. In *Eq. 3*, *z*(*t*) is reset to 0 after an accumulative preset threshold *Z*_threshold_ within a moving window is reached to trigger an escape-type pain behavior (e.g, paw withdrawal). *Equation 4* imposes an inverse sigmoid-shaped relationship between the input amplitude *x*(*t*) and time constant *τ_z_*. The initial pain percept *z*(0) is either zero (no-anticipation), negative value (placebo), or positive value (nocebo or pain anticipation).

**Table 1:** Summary of key results of two computational models and the associated experimental support.

| Condition | Computer modeling results |  | Experimental support |
| --- | --- | --- | --- |
|  | phenomenological model | mean field model |  |
| model description | Eqs. 1-6, Fig. 1D, Table 2 | Eqs. 8-14, Fig. 2B, Table 3 | n/a |
| evoked pain | Figs. 4A-C, Fig. 5A | Fig. 6A-C | Fig. 3C,D, rat ACC and S1 LFP data reported in (Xiao et al., 2019) |
| non-evoked pain | Figs. 4D,E, Fig. 5B | Fig. 6D | Fig. 3B,C,D, rat ACC and S1 LFP data reported in (Xiao et al., 2019) |
| chronic pain | n/a | Fig. 7B,C,H,<br>Fig. 7A,D-H, Fig. 8A-C | pain aversion in rat ACC neurons (Zhang et al., 2017)<br>Fig. 3E,F, rat S1→ACC projection (Singh et al., 2020) |
| placebo & nocebo | n/a | Fig. 8D,E | none found |
| stimulus prediction | n/a | Fig. 9A-C | human S1 activity (Gross et al., 2007; Zhang et al., 2012) |
| pain anticipation | n/a | Fig. 10A<br>Fig. 10B,C | rat's behavior (Urien et al., 2018)<br>rat ACC activity (Urien et al., 2018) |

Notably, *Equations 1* and *2* are reminiscent of modified Kalman filtering operations, where the precision parameter Π_0_ can be viewed as the Kalman gain in the Kalman filtering formulation of predictive coding (Fig. 1*B*). The gain Π_0_ encodes the confidence placed in PEs (Fig. 2*C*) and therefore controls the rate of evidence accumulation or effective step size of the update dynamics of expectation (*Eq. 2*).

**Fig. 2:**
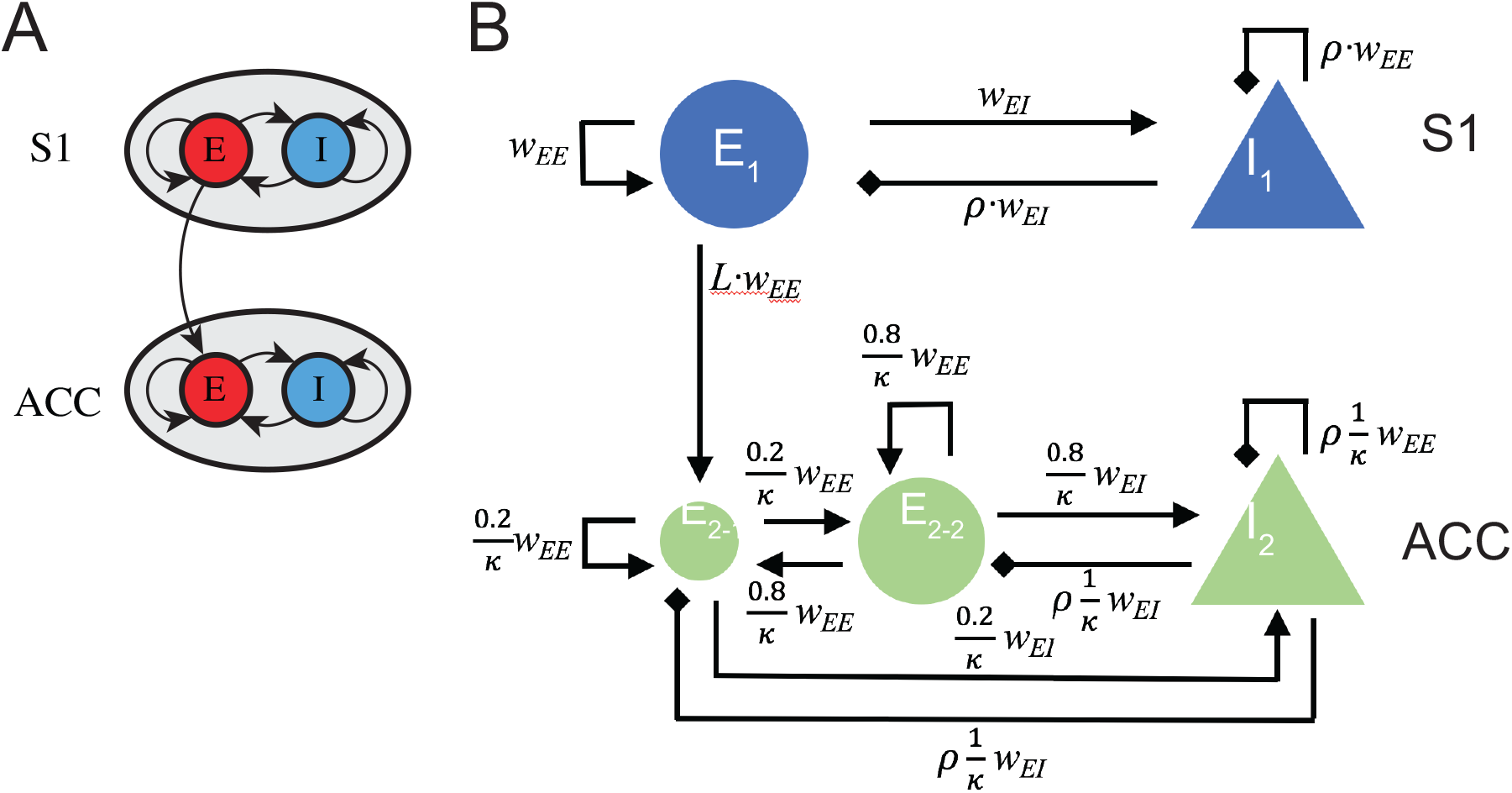
A schematic of mean field model for the S1 and ACC circuits. (*A*) In a reduced model, each brain area is described by an excitatory (E) and an inhibitory (I) population of neurons, with inter- and intra-population coupling. The S1→ACC coupling is assumed to be excitatory and unidirectional. (*B*) A detailed mean-field model that account for biological constraints and details. The pain-responsive ACC neuronal population, *E*_2-1_, is assumed to receive a direct excitatory input from the S1 population *E*_1_. *w_EE_* represents the basic coupling strength between the same type of neuronal populations (E-E or I-I), *w_EI_* represents the basic coupling strength between different types of neuronal populations (E-I or I-E). *ρ* is a negative number that scales the strength of inhibitory input from I-neurons. *L* < 1 is a positive number that scales the effect of long-range S1→ACC projection. *κ* represents the size ratio of S1 population to ACC population.

For the observed neural response variables, in the bottom-up pathway, we assume that dynamics of response variable *u* are driven by the absolute PE as follows

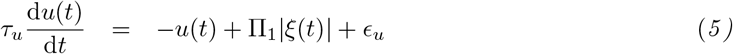

where *τ_u_* > 0 denotes the time constant; *ϵ_u_* denotes the additive Gaussian noise. The gain Π_1_ denotes the precision (inverse of variance) parameter, which weights the *absolute* PE in *Eq*. *5*. We refer to the weighted term Π_1_ × |*ξ*(*t*)| as the “surprise” signal. To see this link, we can assume that there is an expectation uncertainty of *z*(*t*), or equivalently, the PE. Provided that *x*(*t*) is deterministic, then the variance of PE is computed as Var[*ξ*(*t*)] = Var[*z*(*t*)] = 1/Π_1_. If the uncertainty of the expectation is large, the step size will be small (or the update will be conservative); if the uncertainty is low, the update will be more aggressive. In the steady state (i.e., 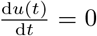), we have *u*(*t*) = Π_1_|*ξ*(*t*)|. Here, the S1 activity encodes the absolute PE or surprise signal, which has been supported by some prior experimental findings (Gross et al., 2007; Arnal & Giraud, 2012; Yu et al., 2019).

The S1 is known to project directly to the ACC (Sesack et al., 1989; Sesack & Pickel, 1992). For the S1→ACC pathway, in the simplest form, we assume that the dynamics of response variable *v* are driven by the signal consisting of a conduction-delayed *u*(*t* − Δ_*u*_) (where Δ_*u*_ > 0) and the pain expectation, as follows

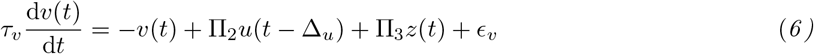

where *τ_v_* > 0 denotes the time constant, and *ϵ_v_* denotes the additive Gaussian noise. Similarly,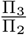defines the relative gain between the two inputs *z*(*t*) and *u*(*t* − Δ_*u*_). The coupling dependency between *u, v* and *z* is shown in Fig. 1*D*.

For convenience, we refer to the model described by *Eqs. 1*-*6* as the *predictive coding model*. The response variables *u* and *v* can be interpreted as the Z-scored LFP gamma and beta power, respectively, which reflect the relative change in the S1 and ACC activity. The choice of conduction time delay Δ_*u*_ reflected the event-related potential (ERP) latency between the S1 and ACC; their time constants *τ_u_* and *τ_v_* were also chosen accordingly. Based on different assumptions of *x* and *z*, we ran computer simulations to produce the dynamics of *u* and *v* from *Eqs. 1*–*6*.

#### Computing time-averaged power

From the simulated traces of *u*(*t*) and *v*(*t*), we computed the averaged power before and after the pain response (e.g., withdrawal). Specifically, let 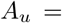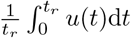 denote the averaged area from the start of computer simulation to the reset (withdrawal) time *t*_*r*_, and let 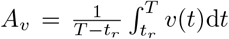 denote the averaged area lasting the same duration it took to reach the *Z*_threshold_ from the reset (withdrawal) time. Therefore, *A_u_* and *A_v_* could be viewed as the averaged pre- and post-withdrawal Z-scored power, respectively. Notably, in the “net” area integration, the curve above 0 contributes to a positive area value, and the curve below 0 contributes to a negative area value.

#### Fourier analysis and spectral asymmetry

Taking the Fourier transform of *Eq. 6* and rearranging the terms, we obtained the mapping of two response variables *u* and *v* in the frequency domain:

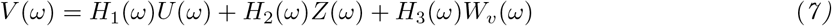

where *H*_3_(*ω*) (or *H*_2_(*ω*)) is a transfer function between *V* (*ω*) and *W_v_*(*ω*)—spectrum for white noise (or *Z*(*ω*), unobserved); and *H*_1_(*ω*) is a transfer function between *V* (*ω*) and *U* (*ω*):

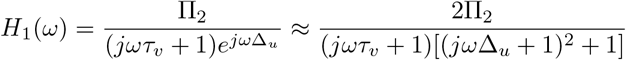

where 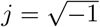, and the approximation is derived from the 2nd-order Taylor series expansion for a small value of *s* (Δ_*u*_ = 0.1 s was used in our computer simulations): 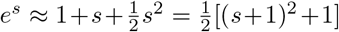. The first term of the denominator in *H*_1_(*ω*) is a 1st-order low-pass filter, and the second term is a 2nd-order low-pass filter. Together, *H*_1_(*ω*) operates as a low-pass filter (Fig. 1*E*) that attenuates the high-frequency (e.g., gamma-band) activity *U* (*ω*), resulting in a lower-frequency (e.g., beta-band) activity *V* (*ω*) in the top-down pathway (Fig. 1*F*). This spectral asymmetry also explains the reason why the Z-scored power is shifted from the S1 gamma-band to the ACC beta-band.

### 2.4 Mean Field Models

To better describe the population neuronal dynamics, we further develop a mechanistic model, with explicit excitatory and inhibitory neuronal populations and synapses, to the predictive coding framework described above. To achieve a trade-off between biological complexity and modeling complexity, we opt for a mean field model (Pinotsis et al., 2014; Wilson & Cowan, 1972).

#### Background

The main assumption of mean field models is that tracking the average activity, such as the mean firing rate and mean synaptic activity, is sufficient when modeling populations of neurons. Given the extensive number of neurons and synapses in even a small area of cortex, this is a reasonable assumption. One of the first mean field models of neural activity is attributed to Wilson and Cowan (Wilson & Cowan, 1972). This two-dimensional model tracks the mean firing rate of an excitatory population of neurons coupled to an inhibitory population of neurons, and has been successfully used to describe visual hallucinations (Ermentrout & Cowan, 1979; Bressloff et al., 2001), binocular rivalry (Wilson et al., 2001), epilepsy (Shusterman & Troy, 2008); Meijer et al., 2015), resting brain state activity (Deco et al., 2011), traveling cortical waves (Wilson et al., 2001; Roberts et al., 2019), and cortical resonant frequencies (Lea-Carnall et al., 2016).

We propose a modified Wilson-Cowan model, with the addition of a synaptic variable for each of the neuronal population. For a single brain area, this amounts to four differential equations (Keeley et al., 2019):

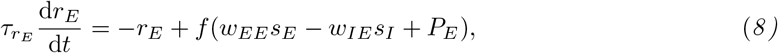

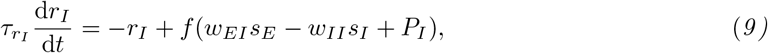

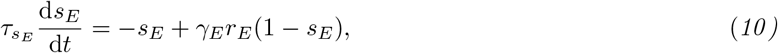

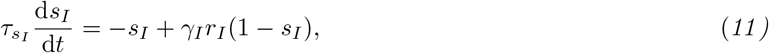

where *r_E/I_* is the population firing rate of the excitatory/inhibitory population, and *s_E/I_* is the synaptic activation of the corresponding population. Each variable has a corresponding time constant *τ*. The inter- and intra-populations coupling strengths are set by {*w_EE_, w_IE_, w_EI_, w_II_*}; *P_E/I_* represents the external input from elsewhere in the cortex; and *γ_E/I_* is the ratio between activation and inactivation times of the synapse. Similar to the standard Wilson-Cowan model, *f* is a sigmoid function:

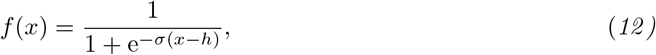

where *σ* is the slope and *h* is the threshold.

In this study, we are interested in the interaction of the S1 and ACC, and consider a model with two excitatory-inhibitory (E-I) pairs, as described by *Eqs. 8*–*11* (Fig. 2*A*). Experimental findings have provided strong evidence that there is a direct S1→ACC projection, which plays an important role in pain processing (Sesack et al., 1989; Sesack & Pickel, 1992; Eto et al., 2011). In contrast, less is known about the role of the ACC→S1 pathway in cortical pain processing. For the sake of simplicity, we first neglected the feedback in our initial model; the impact of feedback will be investigated and discussed later (DISCUSSION).

#### Biologically-constrained mean field model

We have recently combined optogenetics and electrophysiology to dissect the ACC circuit in pain processing (Singh et al., 2020). We have found a direct S1→ACC projection engaged in cortical pain processing. In naive rats, only a small percentage of the ACC population was pain responsive (10-15%). Among those pain responsive neurons, about 20% of the population received a direct input from the S1 (Fig. 3*E*). Among the ACC neurons that receive input from the S1, 37% of them were pain responsive. However in CFA rats, those two percentages increased to 32% and 52%, respectively (Fig. 3*F*).

**Fig. 3:**
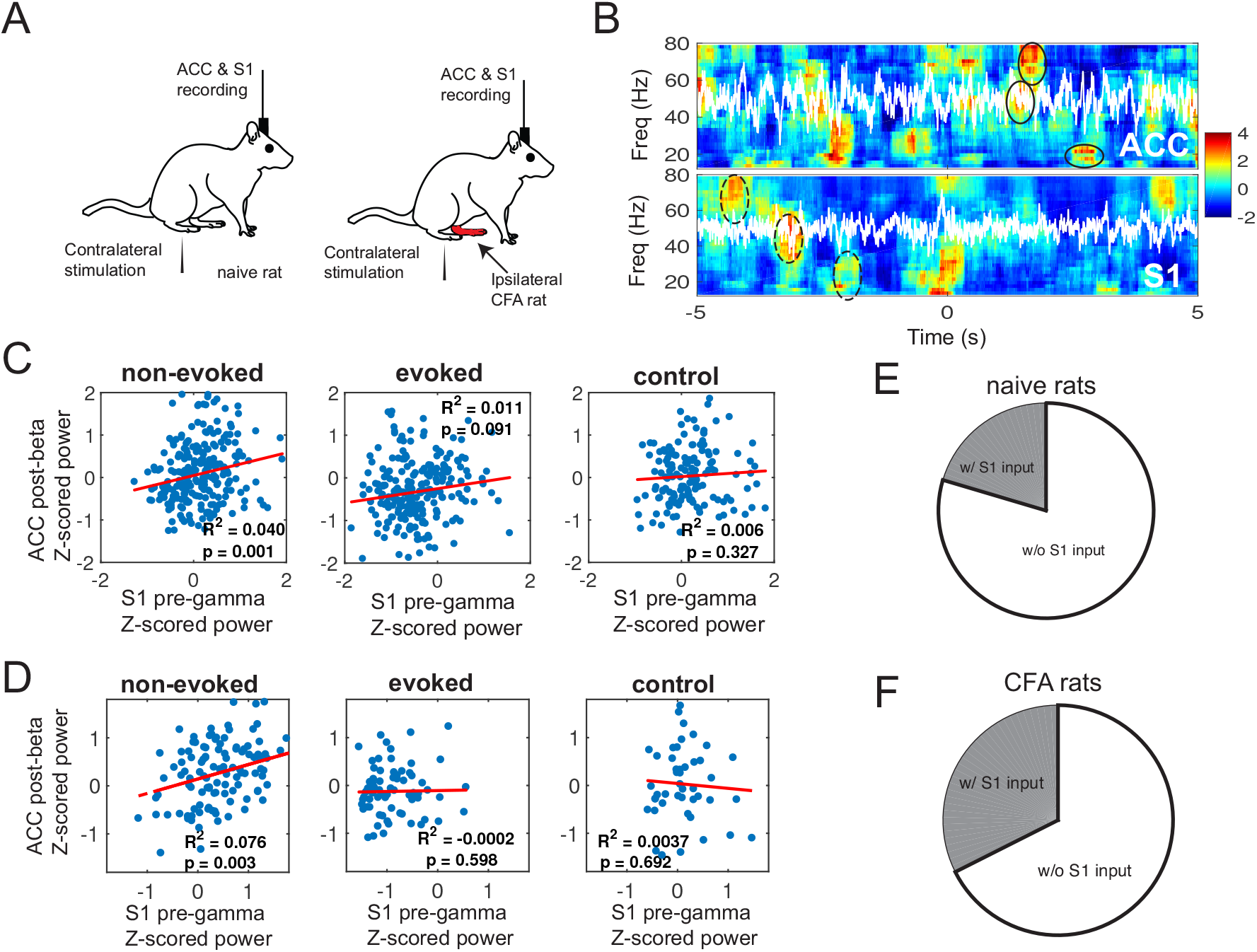
Results excerpted from our previous experimental findings. (*A*) Schematic diagram of noxious stimulation and electrophysiological recording in naive and CFA rats. (*B*) Z-scored spectrograms in the ACC and S1 during a representative non-evoked nociception episode. White traces show the principal component of multichannel LFPs. Time 0 marks the onset of non-evoked nociception event. The post-event power was Z-scored with respect to [−5, 0] s, whereas the pre-event power was Z-scored with respect to [0, 5] s. The S1-ERD during the pre-event period and the ACC-ERS during the post-event period were highlighted by dashed and solid ellipses, respectively. (*C*) Time-averaged Z-scored pre-gamma S1 activity vs. post-beta ACC activity (*n* = 252 non-evoked nociception events, *n* = 233 evoked pain events; *n* = 149 negative controls), for naive rats. In each panel, R-square (i.e., the square of Pearson’s correlation) and *p* values are reported. (*D*) Same as panel C, except for CFA rats (*n* = 127 non-evoked nociception events, *n* = 71 evoked pain events; *n* = 49 negative controls). (*E*) Pie chart of pain-responsive ACC neurons that receive a direct S1 input for naive rats. (*F*) Same as panel *E*, except for CFA rats.

Based on these findings, we made two modifications to the computational model. First, the S1→ACC pathway is modeled with the inclusion of an additional term in *Eq. 8* for the ACC population; namely, we changed the input 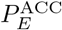 to 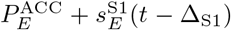, where the excitatory input from the S1 is delayed by a positive Δ_S1_.

Second, we divided the excitatory ACC neuronal population into two subpopulations *E*_2-1_ and *E*_2-2_ (Fig. 2*B*), one of which directly receives S1 input (*E*_2-1_), while the other is indirectly driven by the former one (*E*_2-2_). Therefore, we revised the model described by *Eqs. 8*–*11* with two excitatory-inhibitory (E-I) groups.

We also scaled the inter- and intra-populations coupling strength by the relative population sizes. For example, if the S1 population is twice as large as the ACC population, then the coupling strength of S1→S1 and S1→ACC would be twice as large as those of ACC→S1 and ACC→ACC, respectively. Here we assumed that there are 20% of ACC excitatory neurons that receive S1 inputs; *κ* is the S1/ACC neuronal population size ratio; *ρ* scales the inhibitory/excitatory strength; *L* is the scaling of long-range projection between the two regions. We set *w_EE_* and *w_EI_* as the basic coupling strength, and set other coupling strength with a proper scaling constant (Fig. 2*B*).

Note that the variables that we used previously to describe sensory input and posterior expectations about pain (i.e., *x* and *z*) under predictive coding are now used as exogenous inputs to our neural mass biophysical model. This allows us to handcraft different levels of nociceptive input, and posterior expectations or perceptual representations of the associated pain. Specifically, we assumed that the external inputs are applied equally to the excitatory and inhibitory populations of the S1 and ACC as follows

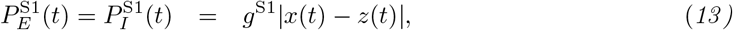

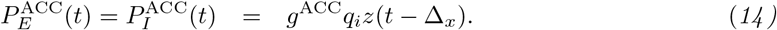

where |*x*(*t*) − *z*(*t*)| denotes the absolute PE, *g*^S1^ and *g*^ACC^ denote two gain parameters for respective neuronal population, and Δ_*x*_ denotes the time delay from the input *x*. Let S1^+^ (or S1^−^) denote the ACC population that receives direct S1 input (or not); let *q_i_* with the subscript index *i* = S1^+^, S1^−^*, I* denote the percentages of pain-responsive neurons in subpopulations *E*_2-1_, *E*_2-2_ and *I*_2_, respectively. The gain parameters of this biophysical model play the same role as the precisions in the predictive coding model.

#### Computing the power using the envelope function

We computed the upper and lower envelopes of the oscillatory firing rate trace. We used the average (midline) of the upper and lower envelopes to calculate the time-averaged synaptic activation variable *s* (or alternatively, the firing rate variable *r*) as a measure of the firing dynamics in our mean field model.

To compute the pre-S1 synaptic activation, we integrated the average power of 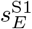 from the baseline (by discarding the initial transient) to the withdrawal onset, and then normalized it by the duration. To compute the post-ACC synaptic activation, we integrated the average power of 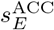 from the withdrawal onset until a fixed window length, and then normalized it by the duration.

#### Software

The custom MATLAB code for implementing two described computational models is distributed online (https://github.com/yuru-eats-celery/pain-coding-model and https://github.com/ymch815/predictive-coding-mean-field-model.git).

## 3 RESULTS

In the following, we first summarize important experimental findings (section 3.1) that were extracted from previous published data (Xiao et al., 2019; Singh et al., 2020), which provide the biological support and motivation for our computational modeling work. Next, we describe our phenomenological model and its simulation results for both evoked pain and non-evoked nociception (section 3.2). We will make data interpretation and prediction related to the experimental results. Finally, we present our computer simulation results based on the mean-field model (section 3.3). In addition to replicating qualitatively similar results as in the phenomenological model, we also adapt the mean-field model for chronic pain and make several experimental predictions. To help the reader understand the materials, we provide a high-level description of these results and connection to experimental findings (Table 1). The rationale and goal of the paper is to motivate the modeling questions based on the empirical experimental findings and make proper interpretations based on the results of model prediction.

### 3.1 S1 and ACC Activity in Naive and Chronic Pain Rats

From the simultaneously recorded S1 and ACC LFP activity, we found that the averaged pre-event Z-scored gamma power in the S1 positively correlated with the averaged post-event Z-scored beta power in the ACC (Fig. 3*C*, left panel). This suggests that pre S1 gamma-ERS (or ERD) was temporally followed by post ACC beta-ERD (or ERS). Notably, the correlated ERS/ERD patterns became weaker during evoked pain episodes (Fig. 3*C*, middle panel) and disappeared in negative control (Fig. 3*C*, right panel). In the chronic pain state of CFA rats, we also found similar observations (Fig. 3*D*).

In our earlier experimental investigation (Singh et al., 2020), we established a direct S1→ACC projection during cortical pain processing. Among pain-responsive ACC neurons, we identified a subpopulation that received the direct S1 input, from both naive and CFA rats (Fig. 3, *E* and *F*, respectively). Compared to naive rats, chronic pain increased the percentage of ACC neurons that received the direct S1 input. Together, these findings provide empirical evidence to characterize chronic pain in our predictive coding model.

In another experimental investigation (Urien et al., 2018), we trained rats with a conditioning paradigm that consists of three experimental phases. During the pre-conditioning phase, we paired a tone (4 kHz, 80 dB, 0.5 s) with a non-noxious thermal stimulus applied to the rat’s hind paw. During the conditioning phase, we paired the same tone with a noxious thermal stimulus to induce pain avoidance. We found that the rat could avoid the noxious stimulus by simply removing its paw after the tone being played, yet before the noxious stimulus being delivered. We also found that a subset of rat ACC neurons responded before the delivery of pain stimulation, and these “pain-anticipating” neurons increased or decreased their firing rates after the tone, and prior to, or in anticipation of, the noxious stimulus. These pain-anticipating neurons gradually shifted their responses to pain and started to respond during the anticipatory period. Later in the post-conditioning phase, these pain-anticipating ACC neurons returned to their baseline behaviors, as the tone stimulus was no longer paired with a noxious stimulus. These data also provide indirect evidence of top-down influence on the ACC neuronal coding.

### 3.2 Computer Simulations for the Predictive Coding Model

The goal of the predictive coding model is to replicate the main findings of the pain experiments at the macroscopic level. From *Eq. 1*-*4*, we ran numerical simulations to characterize the relationship of the surrogate of LFP oscillatory activity between the S1 and ACC. In the following computer simulations, we used the default parameters listed in Table 2. The additive Gaussian noise components {*ϵ_u_, ϵ_v_, ϵ_z_*} were all assumed to have zero mean and unit variance. In each condition, we reported the mean statistics based on 30 independent Monte Carlo simulations, and ran 400 simulations to compute the correlation statistics.

**Table 2:** Summary of default parameters used in the predictive coding model

| Parameter | Value |
| --- | --- |
| $dt$ | 1 ms |
| Time constant $\tau_u$ | 300 ms |
| Time constant $\tau_v$ | 100 ms |
| Time delay $\Delta_u$ | 100 ms |
| Time delay $\Delta_x$ | 300 ms |
| Gain parameter $\Pi_0$ | 1 |
| Gain parameter $\Pi_1$ | 1 |
| Gain parameter $\Pi_2$ | 1 |
| Gain parameter $\Pi_3$ | 1 |
| Threshold $Z_{\text{threshold}}$ | 200 |
| $a$ in Eq. 4 | 5000 |
| $b$ in Eq. 4 | 1 |

To relate our model notations with experimental data, we viewed the variables *u* and *v* as the Z-scored S1 and ACC population neuronal activity (therefore their initial conditions were set to zeros). We also viewed *A_u_* and *A_v_* as the averaged pre- and post-withdrawal Z-scored power from the S1 and ACC, respectively; which corresponded to the S1 LFP pre-gamma Z-scored power and ACC LFP post-beta Z-score power (Fig. 3, *C* and *D*).

#### Evoked pain

In the evoked pain condition, we set the initial pain expectation to be zero (i.e., *z*(0) = 0), and we set *u*(0) = 0 and *v*(0) = 0 for the initial Z-scored activity from the S1 and ACC. In addition, we assumed the finite-duration constant external input, and set the pain stimulus *x*(*t*) = 2 if *t* ∈ [4, 4.5] s and *x*(*t*) = 0 otherwise. The first 4-s period prior to the stimulus was treated as the baseline. Given the initial condition, we ran numerical simulations using the forward Euler method with time step 1 ms. An illustration of representative traces is shown in Fig. 4*A*. As seen in the figure, the *z*-trace closely followed the *x*-trace; the *u*-trace reached an initial peak and gradually decays; and the *v*-trace decayed slower than the other traces.

**Fig. 4:**
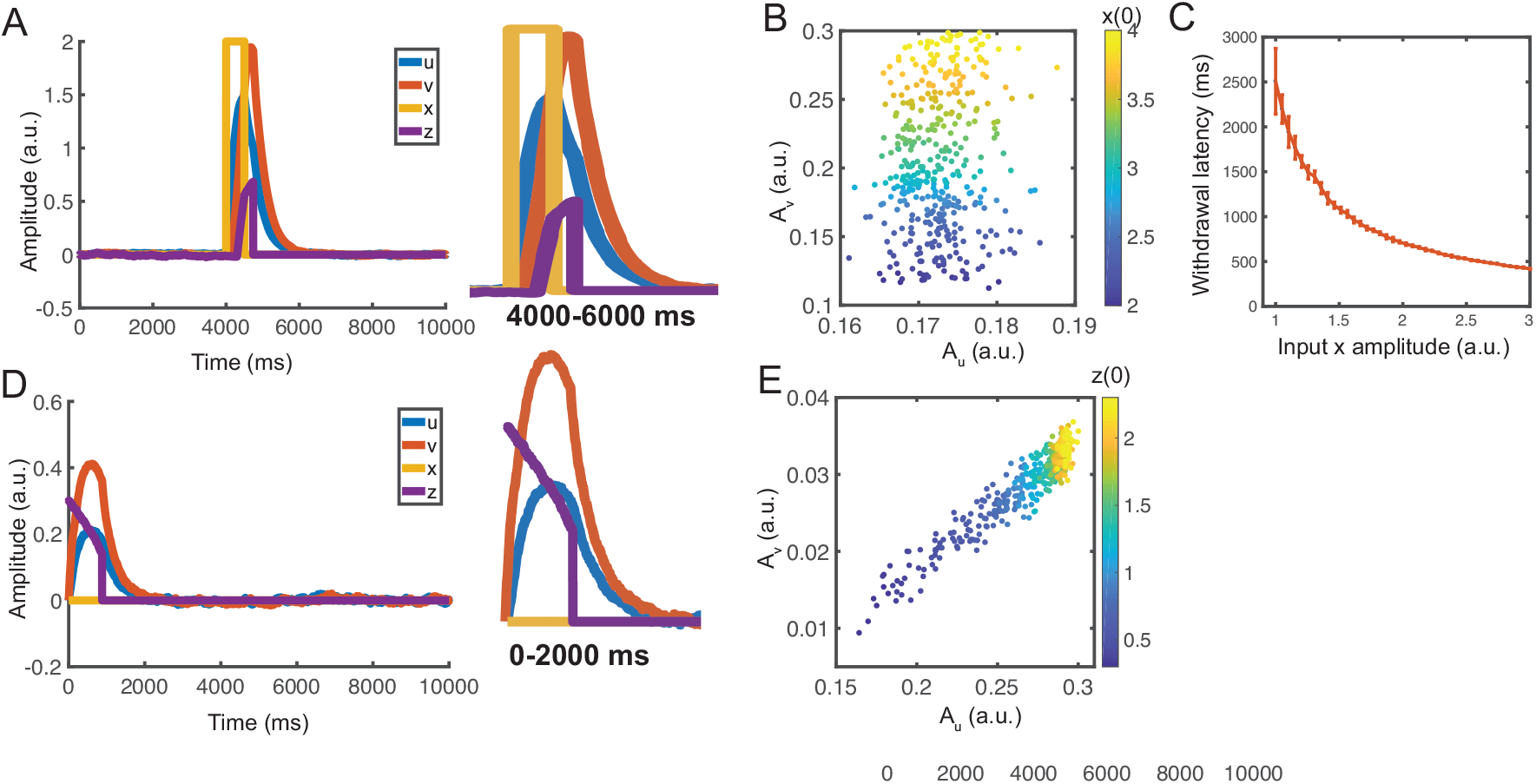
Simulation results from the predictive coding model. (*A*) Simulated 10-s temporal traces of {*x, z, u, v*} in evoked pain. Y-axis is in arbitrary unit (a.u.). Stimulus onset separates the pre-event from post-event periods. Here, we used *u*(0) = *v*(0) = *z*(0) = 0. The right panel shows the zoom-in window of 4000-6000 ms. (*B*) The correlation of *A_u_* (area under *u*-curve) and *A_v_* (area under *v*-curve) was small during evoked pain. Each point was derived from a simulation with a different input amplitude (correlation: 0.097, *p* = 0.06, *n* = 400). The *A_u_* and *A_v_* represent the proxy of average induced responses in the gamma and beta bands. Each point was derived from a simulation with a different *z*(0). (*C*) Reset (withdrawal) latency decreases with increasing input amplitude. Error bar represents the standard error of mean (SEM) (*n* = 50). (*D*) Simulated temporal traces of {*x, z, u, v*} in non-evoked nociception. Y-axis is in a.u. Here, we used *u*(0) = *v*(0) = 0*, z*(0) = 0.3.The right panel shows the zoom-in window of 0-2000 ms. (*E*) *A_u_* (during the pre-event period) was positively correlated with *A_v_* (during the post-event period). Each point was derived from a simulation with a different *z*(0) (correlation: 0.947, *p* < 10^−10^, *n* = 400).

Once *z*-trace was reset upon reaching a threshold, we assumed that moment as the withdrawal onset. We computed the net area under *u*-trace between the start of stimulation to withdrawal (i.e., *A_u_*), as well as the area under *v*-trace between the withdrawal withdrawal and the end of stimulation (i.e., *A_v_*) (Fig. 4*B*).

The withdrawal latency following the stimulus onset is a standard measure to quantify the acute pain behavior (Deuis et al., 2017). In our simulations, we used the duration between the onset of input *x*(*t*) and the time of *z*(*t*) reset as the proxy of withdrawal latency. We found that the latency decreases with increased stimulus intensity or input amplitude (Fig. 4*C*), which is consistent with prior experimental observations (Dirig et al., 1997).

#### Non-evoked nociception

In the non-evoked nociception condition, we set *u*(0) = 0*, v*(0) = 0*, z*(0) = 0.3, *x*(*t*) = 0 (i.e., no stimulus), and *Z*_threshold_ = 200. An illustration of representative traces is shown in Fig. 4*D*. In this example, the *z*-trace decays exponentially until it reached the reset threshold; the *u* and *v*-traces first rose and then decayed exponentially, and it took a longer time for the *v*-trace to approach the baseline.

To introduce trial variability, we assigned *z*(0) with random values. By varying the initial condition *z*(0), we obtained various mean statistics for *A_u_* and *A_v_* during non-evoked nociception. A strong positive correlation between *A_u_* and *A_v_* (Fig. 4*E*) was found. In contrast, the correlation between *A_u_* and *A_v_* was weaker in the simulated evoked pain condition (Fig. 4*B*). These results are consistent with our experimental findings (Fig. 1*B*).

Notably, although these simulations were done in the idealized conditions, and the exact outcome may vary depending on the exact stimulation parameter setup, our computational modeling provides a principled way to investigate the impact of parameters on the read-out phenomenon. We will present such examples below.

#### Sensitivity analysis of gain parameters and transmission delay

Thus far, we have kept all gain (or precision) parameters in unity. Next, we investigated how the change of gain parameter affects the dynamics. Since Var[*z*(0)] = 1/Π_1_, we further assumed that Π_3_ = Π_1_ in *Eq. 6*. To investigate the impact of gain parameters, we considered two scenarios. In the first scenario, we set Π_2_ = 1 and systematically varied Π_1_ and Π_3_ together. In the second scenario, we set Π_1_ = Π_3_ = 1 and systematically varied Π_2_. In both scenarios, the ratio Π_3_/Π_2_ would deviate from unity. The results from these two scenarios are shown in Fig. 5. The qualitative phenomenon that describes the correlation between *A_u_* and *A_v_* was relatively robust with a wide range of gain parameters. In the evoked pain condition, the correlation value remained low. In the non-evoked nociception condition, the correlation value showed an increasing trend with increasing Π_1_ and Π_3_, and showed a decreasing trend with increasing Π_2_.

**Fig. 5:**
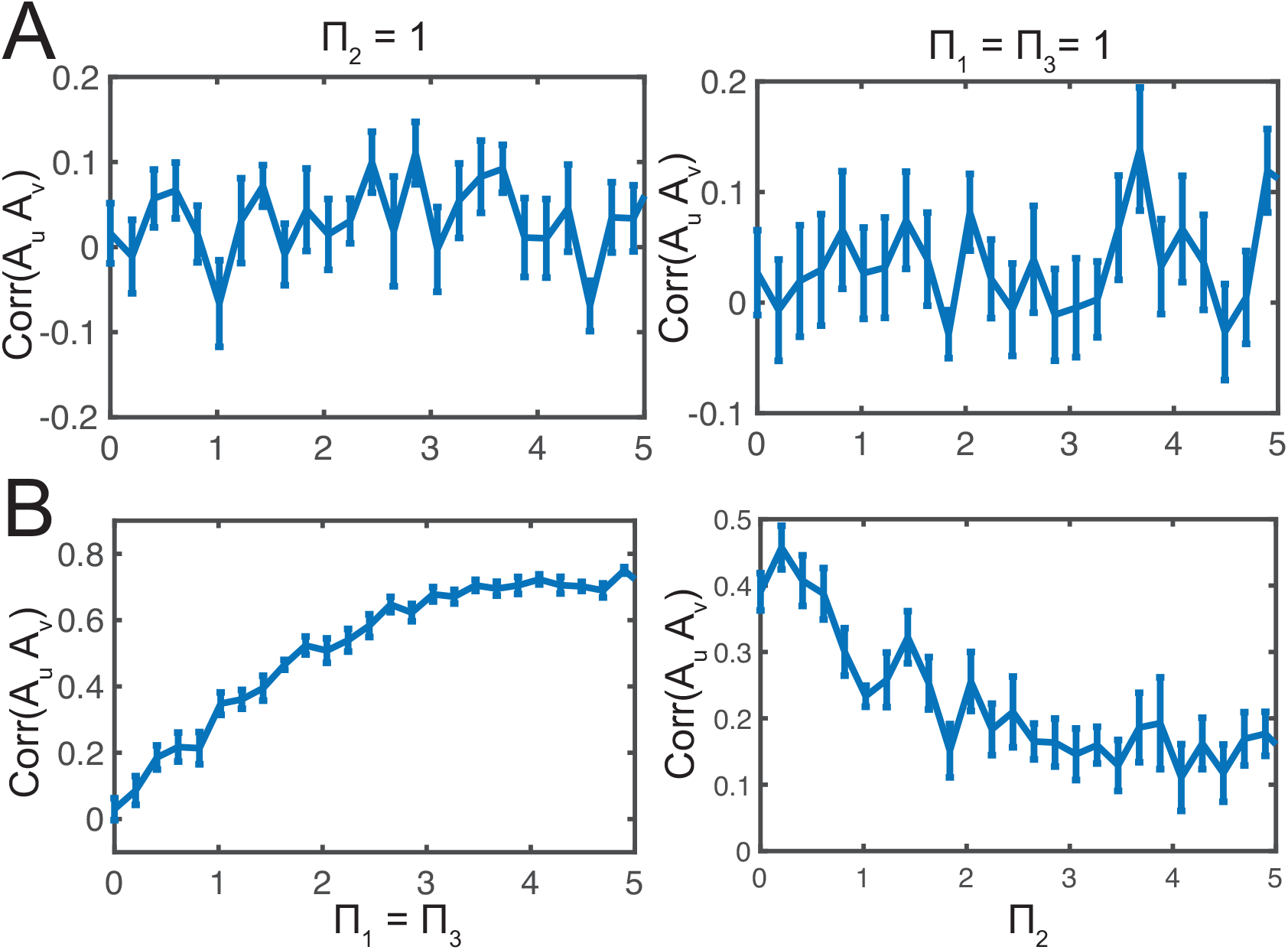
Sensitivity analysis of correlation between *A_u_* and *A_v_* with respect to the gain parameters {Π_1_, Π_2_, Π_3_} in the predictive coding model. (*A*) Evoked pain (*B*) non-evoked nociception. All error bars denote SEM (*n* = 10) and each correlation was computed from 25 trials with random initial conditions.

During task behaviors, the cortico-cortical conduction delay may vary. For cortical communications over long-range connections or information relay between multiple brain areas, the transmission delay may be even longer. To examine the impact of cortico-cortical transmission delay, we further varied Δ_*u*_ and investigated the correlation statistic (Supplementary Fig. 1). We found that the correlation between *A_u_* and *A_v_* was stable for a wide range of delay parameters.

In summary, these simulation results from the predictive coding model replicate several key findings from two experimental pain conditions. However, this phenomenological model is rather abstract, therefore the interpretation of the model parameter or results remains limited. Next, motivated by the neural mass model in the literature (Friston et al., 2012; Bastos et al., 2015), we extended the same line of investigations using the mean field model.

### 3.3 Computer Simulations for the Mean Field Model

In the following computer simulations, we used the default parameters listed in Table 3. We used the forward Euler method to numerically simulate the population dynamics for a total 5.5 seconds (time step 0.1 ms). The pulse input *x* had a 200-ms duration. A 2-s simulation interval was treated as the baseline period. The initial values of all *r_E/I_* and *s_E/I_* were set to zero. We computed the midline envelopes of the synaptic activation variable *s* and firing rate variable *r* of excitatory populations from the S1 and ACC. Since the synaptic activation variable *s* was highly correlated with the firing rate variable *r* (Supplementary Fig. 2), we have used *s* to represent the firing dynamics of the S1 and ACC populations. Notably, the mean field model employs different time constants and delay parameters, as it captures a different spatiotemporal scale from the previous phenomenological model.

**Table 3:** Summary of standard mean field model parameters

| Parameter | Value |
| --- | --- |
| Firing rate of E/I population $r_{E/I}$ | |
| Synaptic activation of E/I population $s_{E/I}$ | |
| Time step in Euler's method | 0.1 ms |
| Percentage of ACC population receiving a direct S1 input | 20% |
| Excitatory synaptic weights $\{w_{EE}, w_{EI}\}$ | 22 |
| Inhibitory synaptic weights $\{w_{II}, w_{IE}\}$ | $w_{II} = \rho w_{EE}$<br>$w_{IE} = \rho w_{EI}$ |
| Scaling parameter $\rho$ for inhibitory/excitatory strength | -1.5 |
| Scaling parameter $L$ for long-range projection | 0.1 |
| S1/ACC population size ratio $\kappa$ | 2 |
| Gain parameter $g^{S1}$ | 2 |
| Gain parameter $g^{ACC}$ | 3 |
| Scaling parameter for % of pain-responsive neurons $q_{S1}^+, q_{S1}^-, q_I$ | 35%, 14%, 10% |
| Slope of sigmoid function $\sigma_E = \sigma_I$ | S1: 0.5; ACC: 0.7 |
| Center of sigmoid function $h_E = h_I$ | S1: 4; ACC: 3 |
| Synaptic activation time constant for excitatory population $\tau_{s,E}$ | 3 ms |
| Synaptic activation time constant for inhibitory population $\tau_{s,I}$ | 10 ms |
| S1 firing time constant for excitatory population $\tau_{r,E}$ | 1 ms |
| S1 firing time constant for inhibitory population $\tau_{r,I}$ | 3 ms |
| ACC firing time constant for excitatory population $\tau_{r,E}$ | 3 ms |
| ACC firing time constant for inhibitory population $\tau_{r,I}$ | 18 ms |
| Ratio between activation and inactivation times of the synapse $\gamma_{E/I}$ | 4 |
| Time delay in S1→ACC projection: $\Delta_{S1}$ | 20 ms |
| Time delay: $\Delta_x$ | 75 ms |

To simulate the withdrawal behavior, the reset time of latent variable *z* was determined when the integration of *z* within a 300-ms moving window reached a predetermined threshold. When *z* was reset to 0, *x* was also simultaneously set to 0, indicating that the animal has escaped from the noxious stimulus. We ran numerical simulations of the mean field model for three pain perception conditions. With different values of *x* and *z*, we simulated the *r_E/I_* and *s_E/I_* dynamics of neuronal subpopulations.

#### Evoked pain and non-evoked nociception

In the evoked pain condition, we set *a* = 2000 (*Eq. 4*), *Z*_threshold_ = 200, and *z*(0) = 0. The dynamics of the populations are shown in Fig. 6*A*. In our simulation, all populations have an oscillatory activity with stable frequency, where the S1 population oscillates in the gamma-band frequency and the ACC population in the beta-band frequency. These oscillatory activities were the emergent property of the local E/I networks as there was no externally imposed stimulus input fed to the network. For each population, we took the upper and lower envelope of oscillation and computed their averaged power as a representation of the mean synaptic activation. As seen in Fig. 6*B*, S1 firing increased quickly after the stimulus onset, as a result of large PE; the activities of two ACC subpopulations increased afterwards, as the latent variable *z* gradually increased. Right after withdrawal, S1 population firing decreased immediately, while ACC population firing decayed slower. Throughout the trial, the ACC subpopulation that received S1 inputs had a greater firing intensity than the ACC subpopulation that did not.

**Fig. 6:**
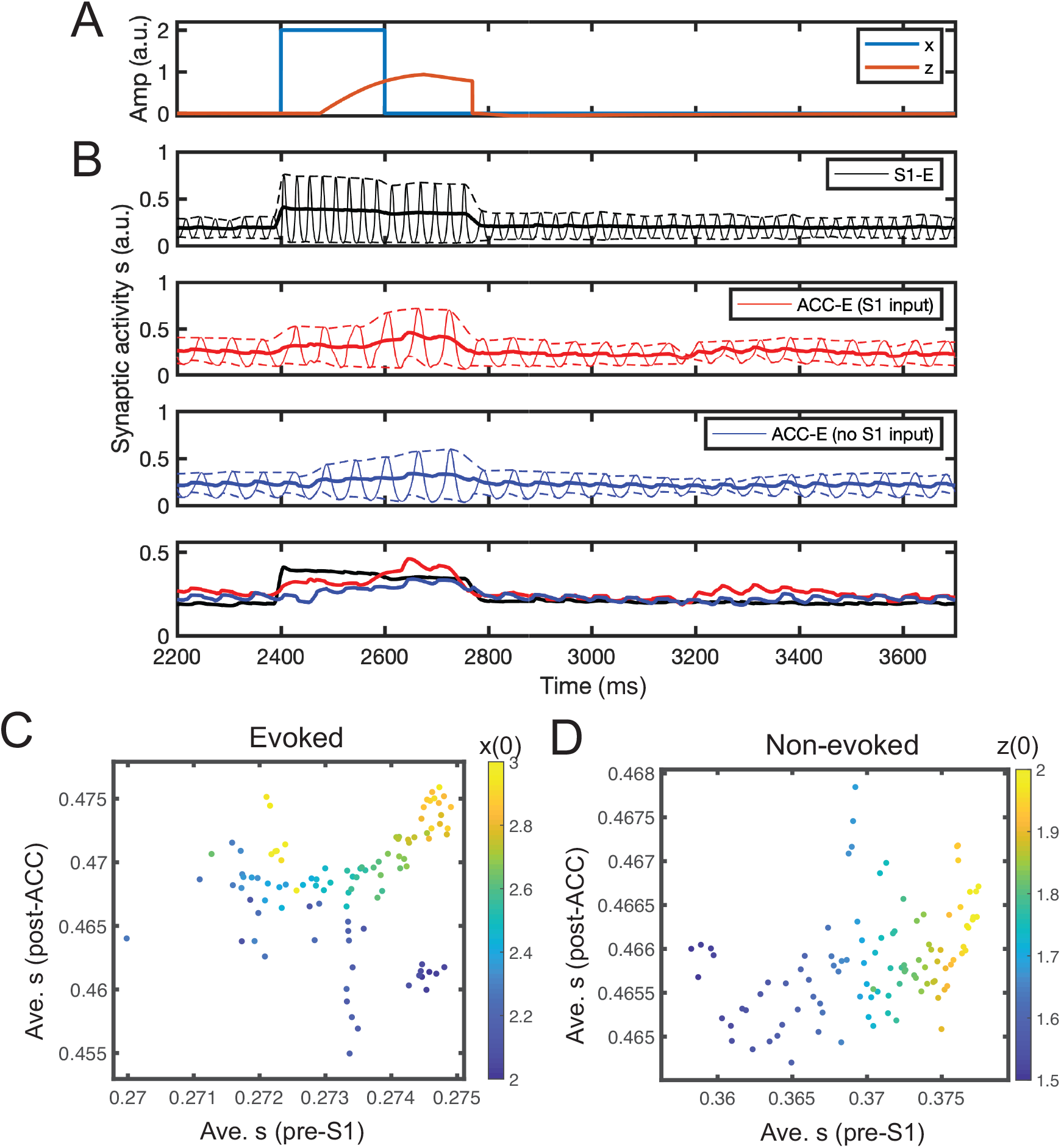
Simulation results from the mean field model. (*A*) Simulated stimulus input *x* and latent *z* trajectories. (*B*) Mean-field activity (synaptic activation *s*) for three different excitatory neuronal populations (*E*_1_*, E*_2-1_*, E*_2-2_) in one evoke pain simulation. Dashed lines show the upper and lower bounds of the envelop around the oscillatory activity. Solid line shows the midline between the upper and lower bounds. For comparison, the last panel replots the three midlines in the first three panels. Time 2400 ms marks the onset time for post-ACC synaptic activation integration. (*C,D*) Scatter plots of average pre-S1 synaptic activation *s* versus average post-ACC synaptic activation *s* derived from the mean field model simulations (*n* = 100) in evoked pain and non-evoked nociception. The Pearson’s correlation coefficients in two panels were 0.15 (*p* = 0.137) and 0.40 (*p* = 4.5 × 10^−5^), respectively. Color bar represents the different initial condition for *x*(0) (panel *C*) or *z*(0) (panel *D*).

We computed time-averaged pre-S1 synaptic activation and post-ACC synaptic activation (METHODS). By varying the stimulus amplitude, we ran 100 Monte Carlo simulations and found that the result was consistent with the previous predictive coding model, showing a relatively weak correlation (Fig. 6*C*).

In the non-evoked nociception condition, we set *Z*_threshold_ = 240 and kept the remaining parameters unchanged. As shown in Supplementary Fig. 3, the firing rates of both S1 and ACC populations increased with a similar pace when the initial top-down expectation *z*(0) was set to a positive value. We computed the pre-S1 and post-ACC activity by varying *z*(0), and found a strong positive correlation between them (Fig. 6*D*), which was again consistent with both experimental findings and the result of previous predictive coding model (Fig. 4*E*).

#### Prediction 1: Placebo condition

Pain perception changes with different contexts. An identical noxious stimulus may cause distinct pain percepts or behaviors depending on the top-down influence. Placebo effects can create real or illusory pain analgesia, which can be pharmacological, psychological, or physical (Wagner & Atlas, 2015). To our best knowledge, there is yet no electrophysiological data available related to the placebo (or nocebo) experiment. Therefore, predictive coding models may be useful to make experimental predictions.

To simulate the placebo effect, we set a negative *z*(0) to represent a biased subjective pain perception. We also set *a* = 2000 and *Z*_threshold_ = 200. The pulse input has a 200-ms duration. As presented in Supplementary Fig. 4, the existence of a negative *z* produced a large PE, driving the S1 population to increase the firing rate, while suppressing the firing of ACC population. After the onset of stimulus, the S1 firing rate increased quickly, while the ACC firing rate increase was slower. By varying *z*(0), we found a positive correlation between the pre-S1 and post-ACC power (Supplementary Fig. 5).

Within the predictive coding framework, placebo-induced treatment expectations can be conceptualized as feedback-mediated predictions, which modulate pain by changing the balance of feedback and feedforward processes at different levels of a neural processing hierarchy (Buchel et al., 2014).

#### Chronic pain

To simulate the chronic pain state, we considered three experimental phenomena observed in chronic pain: (i) the increasing percentage of ACC neurons that receive the S1 input, (ii) the increasing percentage of pain-responsive neurons in each ACC subpopulation, and (iii) the activation of the S1→ACC pathway (Singh et al., 2020). We focused on the targeted ACC neuron population. Specifically, we increased the percentage of ACC neurons that receive direct S1 input from 20% to 30%, increased the scaling parameter *q*_*S*1_+ from 35% to 60%, *q*_*S*1−_ from 14% to 20%, and *q_I_* from 10% to 25%, and increased *L* from 0.1 to 0.2. Other model parameters were kept unchanged (see Fig. 7*A*).

**Fig. 7:**
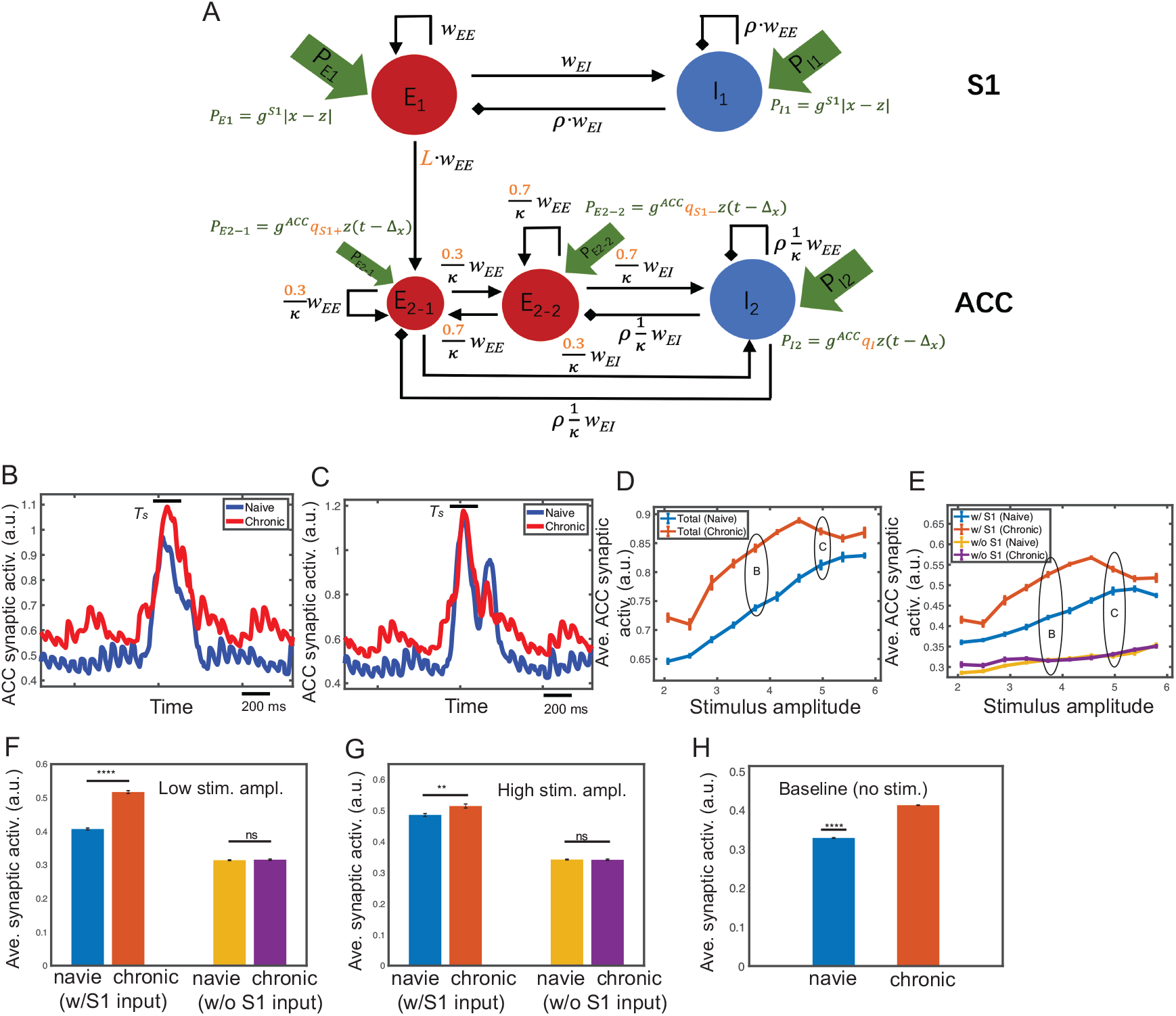
Mean field model simulation results of evoked pain under the chronic pain condition. Fig 7. Mean field model simulation results of evoked pain for the chronic pain condition. (*A*) Modified mean field model (compared to Fig. 3*B*) for chronic pain. Modified variables are marked in red. (*B*) Comparison of the midline envelope of ACC synaptic activation variable *s* between naive (blue) and chronic pain (red) condition for the stimulus amplitude 4.0. Bar above the curve marks the duration *T_s_* between the stimulus onset to withdrawal. (*C*) Same as panel *B*, except for the stimulus amplitude 5.0. In all following plots, the noises are set as *ϵ_z_* = 0.1, *ϵ_E_* = *ϵ_I_* = 0.005. (*D*) Average ACC synaptic activation variable *s* from total population during *T_s_* for varying stimulus amplitude under naive (blue) and chronic pain (red) condition. (*E*) Similar to panel *D*, except for two ACC subpopulations *E*_2-1_ (w/ direct S1 input) and *E*_2-2_ (w/o S1). Mean and SEM for each group are shown. 100 Monte Carlo runs were run with random initial input amplitude *x* ∈ [1.9, 6.0]. (*F,G*) From panel *E*, we replotted the average ACC synaptic activation of *E*_2-1_ (w/ S1) and *E*_2-2_ (w/o S1) during *T_s_* for low and high stimulus amplitude. Error bars were computed from 10 trials with random initial input amplitude *x* ∈ [3.3, 3.7]. For the low stimulus amplitude, there was a significant difference between naive and chronic pain for *E*_2-1_ (*p* < 0.0001, rank-sum test); but not significant for *E*_2-2_ (*p* = 0.86). For the high stimulus amplitude, there was a less significant difference between naive and chronic pain for *E*_2-1_ (*p* = 0.0028); however, the difference of that for *E*_2-2_ was insignificant (*p* = 0.799). (*H*) Average ACC synaptic activation during baseline (no stimulus) was significantly lower (*p* < 0.0001, rank-sum test) in naive (blue) than in chronic pain (red). Average was computed from the baseline ([0.5, 1.5] s) by discarding the initial transient period.

In the evoked pain condition, we first computed the traces of synaptic variable *s* in single-trial simulations. As expected, with a relatively low stimulus (*x* = 4.0), the ACC population had a significantly higher firing intensity in the chronic pain condition than in the naive case (Fig. 7*B*). When the stimulus was sufficiently high (*x* = 5.0), the ACC population had a similar firing intensity in both chronic and naive situations (Fig. 7*C*).

Next, by varying the stimulus amplitude from 1.9 to 6.0, we ran 100 trials and computed the averaged synaptic variable *s* of the ACC population from the stimulus onset to withdrawal, which reflects the overall firing intensity of ACC neurons in response to the stimulus. As shown in Fig. 7*D*, the difference in firing intensity between naive and chronic pain conditions increased with increasing stimulus amplitude in the presence of low-intensity stimulus; whereas this difference diminished with increasing stimulus amplitude in the presence of high-intensity stimulus. With chronic pain, the ACC firing intensity increased disproportionally depending on the stimulus intensity, which is consistent with our previous experimental findings (Zhang et al., 2017). In addition, we have computed the maximum synaptic activation as well as the latency to the maximum (Supplementary Fig. 6).

We then considered the activities of ACC subpopulations *E*_2-1_ and *E*_2-2_ separately. As shown in Fig. 7*E*, with a low-intensity stimulus, the response of the ACC subpopulation with the S1 input was similar to the response of total population, showing a significant increase in the firing rate from naive to chronic pain. However, the ACC subpopulation without receiving the S1 input did not change their firing rate significantly (Fig. 7*F*). This suggested that the disproportional increase in ACC firing intensity from naive to chronic pain was contributed mainly by neurons that received the S1 input. With a high-intensity stimulus, the difference in the firing rate between naive and chronic pain conditions was small for the ACC subpopulation with the S1 input, but the difference was still not significant for the ACC subpopulation without the S1 input (Fig. 7*G*).

Experimentally, the ACC baseline firing rate was higher in chronic pain than the naive condition (Singh et al., 2020). Our model prediction also supported this result (Fig. 7*H*), where the time-average of ACC baseline activity was computed over the period [0.5, 1.5] s.

In the absence of stimulus, we found that chronic pain induced more sustainable high firing intensity in the ACC. We computed the fraction of time when *s* was greater than a certain threshold within the period *T_s_* (from *z* onset to withdrawal), and found a sigmoid-like shape with increasing *z*(0) (Fig. 8*C*). From naive to chronic pain, the sigmoid curve shifted toward the left, which indicated that the fraction of time saturated at a lower *z* level in the chronic pain condition. As shown in Fig. 8*A*, when *z* was low, chronic pain induced a higher and sustainable firing response compared with the naive condition. In contrast, when *z* was high, both curves were saturated so that the time above threshold was nearly the same in both conditions (Fig. 8*B*). This implies that if spontaneous pain (i.e., non-evoked nociception) was primarily induced by a top-down input, then the nociceptive response of ACC neurons would be more sustainable in the chronic pain condition than in the naive condition.

**Fig. 8:**
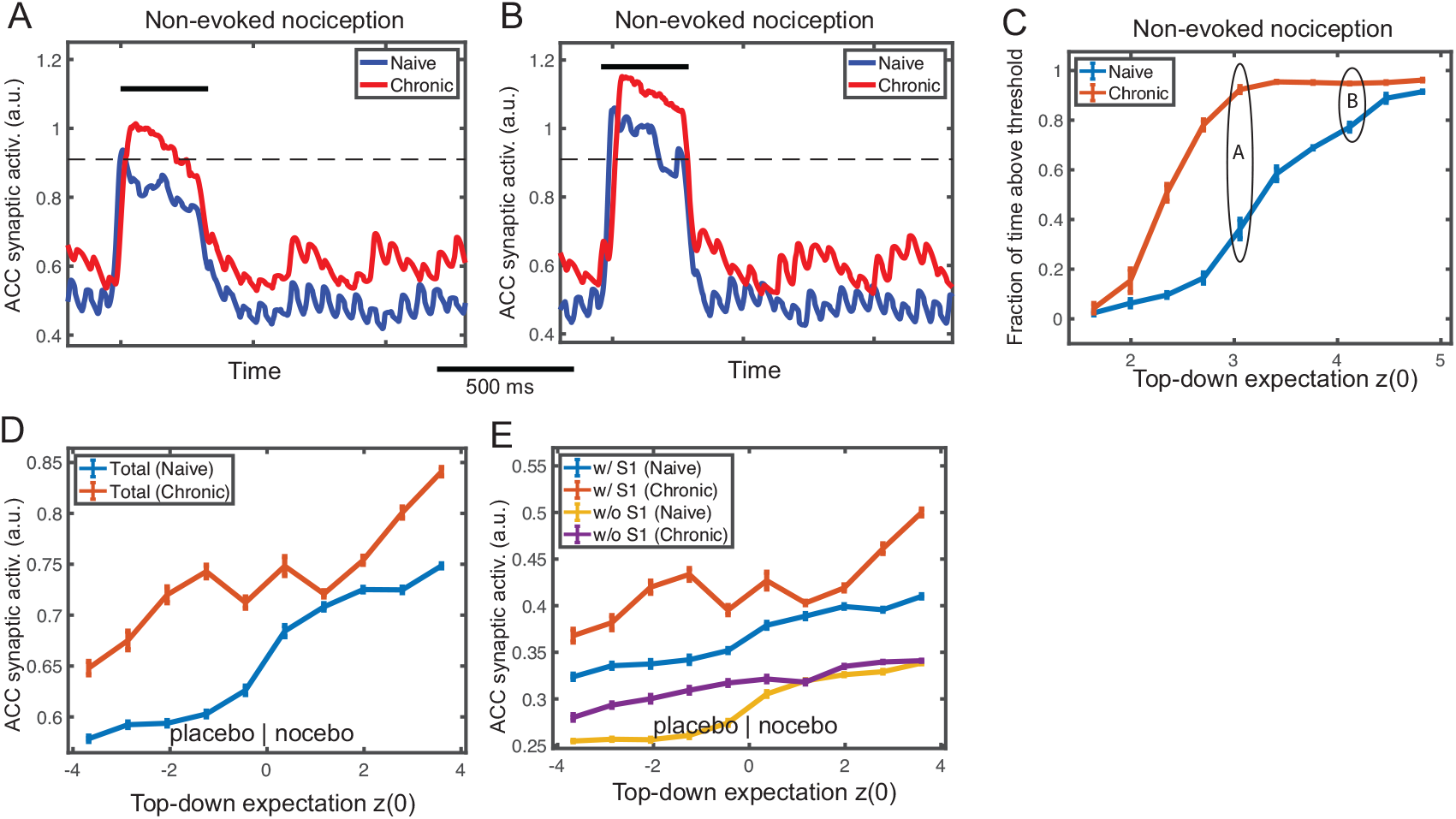
Mean field model simulation results of non-evoked nociception and placebo/nocebo effects under the chronic pain condition. (*A*) Simulated midline envelope trace of ACC synaptic activation variable *s* in non-evoked nociception under naive (blue) and chronic pain (red) conditions, for an initial top-down expectation *z*(0) = 3.0. (*B*) Similar to panel *A*, except for *z*(0) = 4.5. At low *z*(0), the fraction of time above threshold during *T_s_* (between the stimulus onset and withdrawal) was longer in the chronic pain condition; at high *z*(0), the fraction was similar between the two conditions. (*C*) Fraction of time during *T_s_* that ACC synaptic activation variable was above the threshold (horizontal dashed line) for various top-down expectation *z*(0) in naive (blue) and chronic pain (red) conditions. The curve has a sigmoidal shape and shifts leftward from naive to chronic pain condition. 100 Monte Carlo trials were run with random *z*(0) ∈ [1.5, 5.0]. Mean and SEM for each group are plotted. (*D*) Comparison of average ACC synaptic activation in placebo/nocebo effects under naive (blue) and chronic pain (red) conditions for various initial top-down expectation *z*(0), where *z*(0) < 0 and *z*(0) > 0 represent the placebo and nocebo effects, respectively. The synaptic activation *s* of total ACC population increased monotonically, and shifted upward from the naive to chronic pain condition. (*E*) Similar to panel *D*, except for ACC subpopulations *E*_2-1_ (w/ S1 input) and *E*_2-2_ (w/o S1 input). The subpopulation *E*_2-1_ had a similar shape of the total population, while *E*_2-2_ did not increase much from the naive to chronic pain condition. 100 Monte Carlo trials were run with random *z*(0) ∈ [−4.0, 4.0]. Mean and SEM are plotted for each group.

In the placebo and nocebo conditions, we predicted a monotonically increasing trend in ACC firing with respect to increasing *z* in both naive and chronic pain states (Fig. 8*D*), where negative *z*(0) corresponded to the placebo effect and positive *z*(0) to the nocebo effect. This is consistent with the definitions of placebo effect as reduced nociceptive responses and the nocebo effect as increased responses. In our simulations, we also found that the curve shifted upward from the naive to chronic pain condition, indicating that the placebo effect was weaker (i.e., feeling less relieved) and the nocebo effect was stronger (i.e., feeling more painful) in chronic pain. The mean firing curve of each subpopulation with regard to *z*(0) (Fig. 8*E*) suggested that the ACC subpopulation receiving the S1 input contributed predominantly to this shift.

#### Precise noxious stimulus prediction decreases the S1 response

Next, we made predictions of the S1 response within the predictive coding framework. Specifically, we fixed the stimulus input *x* and investigate how the S1 firing intensity would change with respect to different *z*(0). As shown in Fig. 9*A*, the pre-stimulus S1 firing intensity increased monotonically with *z*(0) However, the post-stimulus S1 firing intensity was determined by the absolute PE, or |*x* − *z*(0)|. As shown in Fig. 9*B*, the curve of post-stimulus S1 firing intensity had a V-shape with respect to *z*(0), and the minimum of V-shape curve shifted rightward when we increased the stimulus amplitude *x*.

**Fig. 9:**
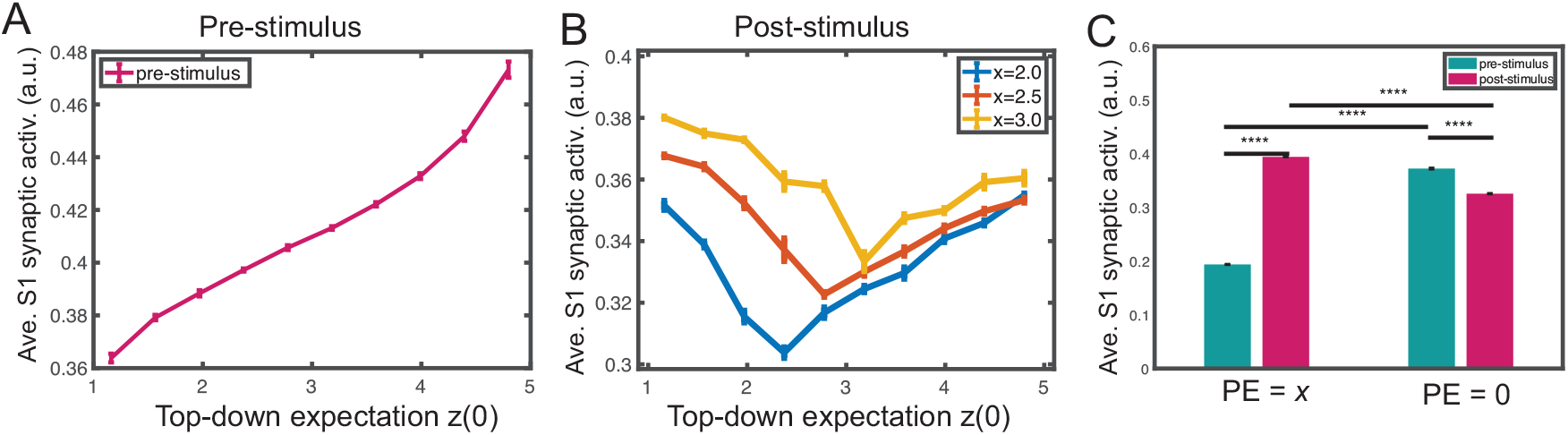
Mean field model simulation results of the S1 synaptic activation variable with regards to the prediction error (PE) and stimulus prediction. (*A*) During pre-stimulus baseline, time-averaged S1 synaptic activation variable increased monotonically with *z*(0). (*B*) During post-stimulus presentation, the average S1 synaptic activation variable exhibited a V-shaped profile from varying stimulus amplitude (*x* = 2.0, 2.5, 3.0), where the minimum occurs when *x* = *z*(0) or PE = 0. The minimum shifted rightward with increasing *x*, indicating that the post-stimulus S1 synaptic activation variable was proportional to |*x* − *z*|. 100 Monte Carlo trials were run with random *z*(0) ∈ [1.0, 5.0]. Mean and SEM for each group are plotted. (*C*) Comparison of average S1 synaptic activation at different time (pre vs. post-stimulus) and PE: PE = *x* (i.e., *z* = 0) and PE = 0 (i.e., *z* = *x*). Ten Monte Carlo trials were run with random input amplitude *x* ∈ [2.0, 2.4]. There was a significant difference in the average S1 synaptic activation between the pre vs. post-stimulus period in both cases. All *p*-values for pair comparisons marked in the graph were less than 0.0001 (rank-sum test). The pre-stimulus firing was computed from the expectation *z* onset (from time 0 if no expectation) to the stimulus *x* onset; the post-stimulus firing was computed from the stimulus onset to withdrawal.

We further examined the effect of prediction on the S1 firing intensity. We set a positive *z*(0) and assumed that *z* remained constant before the stimulus onset, which represented a prediction of the stimulus *x*. We tested two scenarios: one with zero PE (i.e., *z*(0) = *x*), the other with a PE of *x* (i.e., *z*(0) = 0). We measured the firing intensity of pre- and post-stimulus S1 excitatory population, respectively. As illustrated in Fig. 9*C*, when PE was *x*, the pre-stimulus S1 firing was significantly lower than the post-stimulus S1 firing; however, the trend was reversed when there was a precise prediction (i.e. PE=0).

It is noteworthy that our model prediction is in line with several experimental findings in the literature. First, human S1 gamma oscillations can predict subjective pain intensity (but not objective stimulus intensity) (Gross et al., 2007; Zhang et al., 2012). Second, the precise prediction of pain stimulus intensity decreases the S1 gamma-band activity (Arnal & Giraud, 2012). Third, the prediction level is positively correlated with the “rating” of pain stimulus (Peng et al., 2015).

#### Pain anticipation shifts the onset of the ACC response

Pain anticipation shifts the onset of the ACC response. Furthermore, we made predictions of the ACC response in the presence of pain anticipation within the predictive coding framework. Based on our prior experimental findings (Urien et al., 2018), we conducted a computer simulation of tone-conditioning pain anticipation (or prediction) experiment. Specifically, the latency of peak firing rate (Fig. 10*A*) of ACC neuronal populations changed significantly in the presence of anticipation or prediction. In the absence of prediction, we observed a positive peak latency, which implies that it took time for the ACC firing rate to accumulate upon the nociceptive stimulus. However, when we set the latent variable *z* to a value that is equal to the stimulus amplitude, the peak ACC firing rate appeared earlier or before the arrival of the actual stimulus—thereby leading to a negative peak latency. This is consistent with what we observed in the rat experiments. Furthermore, we tested the mean and peak firing activity during the 50-ms tone period right after we set *z* to the stimulus amplitude (Fig. 10*B* and Fig. 10*C*). In the case of pain anticipation, both the mean and peak firing rates during the tone-conditioning period increased significantly compared to those in the absence of anticipation. This result indicated that the existence of anticipation was correlated with the activation of ACC neurons during the tone period. Therefore, our computer simulations can replicate the experimental findings of ACC neurons.

**Fig. 10:**
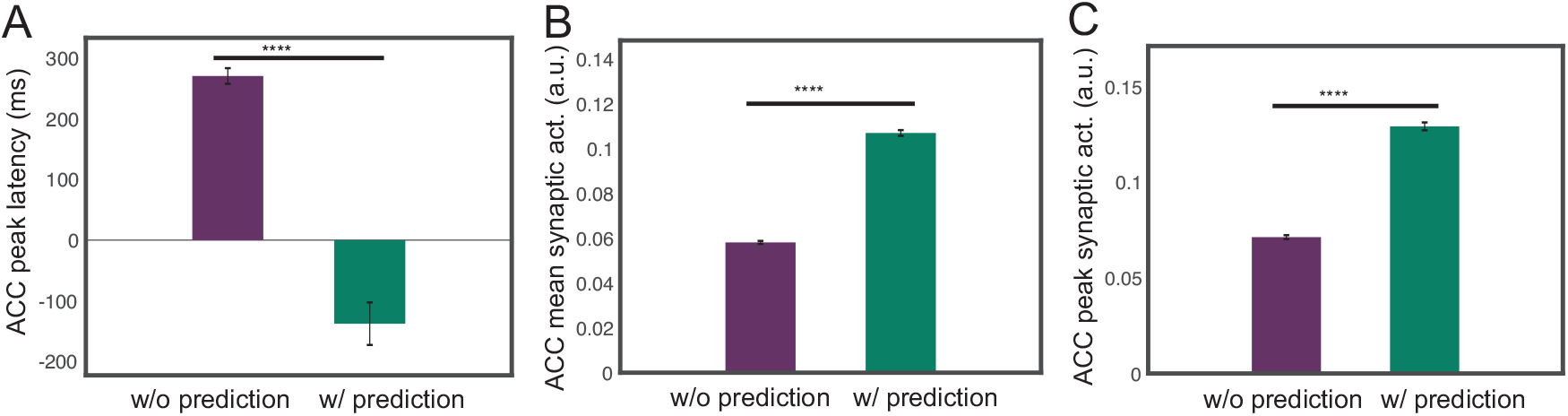
Mean field model results of the ACC firing activity in a simulated tone-conditioning pain anticipation experiment. Ten Monte Carlo trials were run with random stimulus input amplitude *x* ∈ [2.0, 2.4]. (*A*) Comparison of the latency of ACC peak synaptic activation with respect to the onset of the stimulus between without prediction and with prediction conditions. ****: *p* < 0.0001 (rank-sum test).(*B*) Comparison of the mean of ACC synaptic activation during the 50-ms tone period between without prediction and with prediction conditions. (*C*) Comparison of the maximum of ACC synaptic activation during 50-ms tone period between no prediction and with prediction conditions.

#### Prediction 2: Influence of ACC→S1 feedback

Thus far, we have only assumed the direct S1→ACC projection in the circuit model based on the available experimental literature (Sesack et al., 1989; Sesack & Pickel, 1992; Eto et al., 2011; Singh et al., 2020). We further asked whether the presence of ACC→S1 feedback changes the model prediction. To explore that answer, we tried incorporating the ACC→S1 feedback into the mean field model, and found qualitatively similar observations in the average S1 synaptic activation variable as the default setup without feedback (Supplementary Fig. 7). This result suggest that if there is an indirect pathway that the ACC activity affect the S1 response, the simulation results of our biophysical model remain approximately valid.

## 4 DISCUSSION

In this work, we have used computational (predictive coding) models and neural mass (biophysical) models to reproduce the same empirical findings (i.e., dissociations in terms of gamma and beta-band neural responses). We accomplished this by choosing model parameters that reproduced the basic findings in terms of pre-and post-pain induced LFP responses observed empirically in rodent experiments. By incorporating biophysical constraints, the neural mass model could well explain the findings in chronic pain.

### 4.1 Neural Pathways for Pain Perception

Limited by rodent neurophysiological recordings, we have only focused our attention on the S1 and ACC circuits in the context of predictive coding. In reality, however, many other cortical or subcortical circuits are also engaged in pain processing. In the ascending (“bottom-up”) pathway, the nociception originates in the peripheral system, and then passes the signals to the dorsal horn of the spinal cord, then further to the thalamus, and the cortex. The descending (“top-down”) system, involving the midbrain, RVM (rostral ventromedial medulla), PAG (periaqueductal gray) and other areas, can exert both inhibitory and modulatory influence. There may be multiple routes of descending control. It may originate in the cortex, including the ACC, and project to the PAG. The PAG in turn sends projections to RVM and then to the spinal cord (Buchel et al., 2014).

The S1 receives the bottom-up sensory input, involving the regulation of cortical excitability. However, the prestimulus S1 gamma oscillations can predict subjective pain intensity (Gross et al., 2007), whereas the precise prediction of pain stimulus intensity decreases the gamma-band activity (Arnal & Giraud, 2012). Together, these results also suggest that the S1 activity can represent the relative mismatch of expectations and sensory evidence (Bauer et al., 2014).

Several lines of experimental evidence have pointed to a direct S1→ACC projection in cortical pain processing (Sesack et al., 1989; Sesack & Pickel, 1992; Eto et al., 2011). We have recently used experimental techniques to establish a direct S1→ACC projection in rats during the course of cortical pain processing. Activation of S1 axon terminals in the ACC can recruit new ACC neurons to respond to noxious stimuli, as well as increase the spiking rates of individual pain-responsive neurons; in the chronic pain state, the S1→ACC connectivity is enhanced, as manifested by a higher percentage of ACC neurons that respond to S1 inputs (Singh et al., 2020). To date, however, it remains unknown whether there is an indirect ACC→S1 pathway through the cortico-cortical feedback loop that modulate pain processing. More experimental investigations are still required in the future.

The ACC is a major target of midbrain dopamine neurons, which encode reward-related information (such as the reward prediction error). The ACC is reciprocally connected with the amygdala and the orbitofrontal cortex (OFC), which has a projection to the nucleus accumbens (NAc). Importantly, the ACC is also reciprocally connected with the prefrontal cortex (PFC), a region implicated in executive control, working memory, and rule learning. Therefore, the ACC may serve as a gateway for incorporating reward-related information into sensorimotor mappings subserved by the PFC (Hayden & Platt, 2009). Moreover, the PFC has been confirmed to play a modulatory role of gain control in pain processing (Dale et al., 2018). Activation of the PFC provides effective relief to sensory and affective pain symptoms via descending projections in rodents (Lee et al., 2015; Martinez et al., 2017; Zhang et al, 2015; Hardy, 1985). Nevertheless, a complete circuit dissection of cortical pain processing between sensory cortices, ACC, OFC, and PFC has not been established.

### 4.2 Experimental Evidence for Predictive Coding in Pain Perception

In the presence of uncertainties, the brain uses a prediction strategy to guide decision making or perceptual inference (Tabor et al., 2017). Within the predictive coding theory, oscillatory beta-band activity has been linked to top-down prediction signals and gamma-band activity to bottom-up PEs (Pelt et al., 2016). Specifically, in a human MEG study, Granger-causal connectivity in the beta-band was found to be strongest for backward top-down connections, whereas the gamma-band was found to be strongest for feed-forward bottom-up connections (Pelt et al., 2016). In our recent Granger causality analysis of rodent S1-ACC LFP data (Guo et al., 2020), we have observed a S1→ACC Granger-causality peak at a higher frequency (~75 Hz), and an ACC→S1 Granger-causality peak at a lower frequency (~55 Hz), supporting this predictive coding theory. This causality analysis result may also be ascribed to the spectral asymmetry in predictive coding (Bastos et al., 2015).

Predictive coding may provide the key to understanding important phenomena in pain perception (Wiech, 2016; Ploner et al., 2017; Morrison et al., 2013). Unlike evoked pain, spontaneous pain or non-evoked nociception is detached from an overt stimulus and may be driven by internal processing inside the pain matrix. An important finding from our previous experimental data (Xiao et al., 2019) is the fact that the pre-S1 gamma-ERD/ERS correlates with the post-ACC beta-ERS/ERD during non-evoked nociception, whereas the correlation becomes weaker or diminishes in evoked pain or baseline (Fig. 1*C,D*). This phenomenon holds in both naive and chronic pain-treated rats, suggesting an information flow between the bottom-up (gamma) and top-down (beta) loops. Therefore, the brain may use differential neuronal responses to represent bottom-up and top-down modulations of pain, and to provide complementary information about pain perception (Tiemann et al. 2015).

The temporal coordination of beta-ERS/ERD and gamma-ERD/ERS between the ACC and S1 during pain perception corroborates with some previous gamma-ERS and alpha or beta-ERD reports on human EEG findings (Gross et al., 2007; Hu et al., 2013; Schultz et al., 2015). The pain-induced alpha or beta-ERS/ERD is highly dependent on the cortical region, which may be related to sensory gating and functional inhibition, or influenced by top-down attention modulation (Peng et al., 2015). In addition, it has been reported that pre-stimulus human EEG oscillations at the alpha (at bilateral central regions) and gamma (at parietal regions) bands negatively modulated the perception of subsequent nociceptive stimuli (Tu et al., 2016).

### 4.3 Chronic Pain

In the chronic pain state, sensory hypersensitivity and aversion are commonly observed. Chronic pain can also alter acute pain intensity representations of noxious stimuli in the ACC to induce generalized enhancement of aversion (Zhang et al., 2017). While cortical pain responses differ between naive and chronic pain animals, the exact mechanisms of transitioning from acute to chronic pain is still incompletely understood. Our computational model can provide valuable predictions to confirm the experimental findings. Our recent experimental data have shown an increased number of ACC neurons that receive S1 nociceptive inputs, and these neurons that receive S1 inputs also have elevated firing rates (Singh et al., 2020).

In chronic pain experiments, CFA mice with inflammatory pain show elevated resting gamma and alpha activity and increased gamma power in response to sub-threshold stimuli, in association with nociceptive hypersensitivity. Inducing gamma oscillations via optogenetic activation of parvalbumin-expressing inhibitory interneurons in the S1 enhances nociceptive sensitivity and induces aversive avoidance behavior (Tan et al., 2019). In addition, the magnitude of placebo analgesia effect appears to be stronger in chronic pain patients experiencing hyperalgesic states (Vase et al., 2014). Our computer simulation results have indirectly supported these findings (Fig. 8 and Fig. 9, respectively).

### 4.4 Limitations

Our have computational models have succeeded in modeling several key experimental data findings (Table 1), including the LFP spectral asymmetry in the S1 and ACC, animal behaviors in evoked pain and pain anticipation, coordinated S1-ACC activity during chronic pain, the S1 activity during stimulus prediction, the ACC activity during pain anticipation.

However, there are also several conceptual limitations in our computational models. First, we did not explicitly model the cortical layer-specific role in the S1 and ACC. It is well known that different cortical layers receive distinct sources of feedforward or feedback input and may carry different computational roles in predictive coding. Specifically, L4 neurons may receive inputs from the thalamic projection; L2/3 pyramidal neurons are critical for receiving prediction signals from high-level cortical areas, and interlaminar connections may support the temporal integration of feedforward inputs and feedback signals to predict future perception (Constantinople & Bruno, 2013; Bastos et al., 2020). Recent fMRI experiments also suggest the predictive coding in the human S1 in a layer-specific manner (Yu et al., 2019). Second, our biophysical models were established based on oversimplified assumptions and have ignored many details in the canonical microcircuit, such as the cell type specificity, thalamic feedback, and neuromodulatory input. The dynamic causal model (DCM) can potentially capture more functional and anatomical properties of the microcircuits for predictive coding (Bastos et al., 2015). However, detailed causal modeling of cortical connectivity is highly challenging (involving many parameters), which is difficult to fit based on rodent LFP recordings alone. Finally, thus far we have only developed mathematical equations to characterize the neural response variables *u*(*t*) and *v*(*t*); in other words, our models are purely phenomenological and descriptive. A computational strategy would be developing practical algorithms to predict the latent *z*(*t*) based on the observed responses {*u*(*t*)*, v*(*t*)}; this will be the subject of our future research. Overall, a computational model is only as good as its assumptions. Although our model predictions depend on the model oversimplification and parameters, the predictive coding modeling framework is sufficiently flexible and powerful to generate rich neuronal population dynamics.

In summary, motivated by empirical experimental findings in rodents, we have developed a predictive coding framework and computational models to characterize the neuronal population activity of the rat S1 and ACC in pain conditions. To our knowledge, our work represents the first effort along this direction. Our first model is phenomenological and characterizes the macroscopic neural activity, whereas the biophysically-constrained mean field model characterizes the mesoscopic neuronal population activity. Importantly, our mean field model imposes biological constrains onto the E/I populations. Our computational models have not only presented a good prediction of the rodent data, but also made experimental predictions on the placebo/nocebo effects; the next step is to further validate the predictive coding models in human pain experiments. This effort would require the use of source localization techniques to reconstruct the S1 and ACC activity based on high-density EEG or MEG recordings (Pelt et al., 2016; Hauck et al., 2015; Zhang et al., 2016). In addition, our computational model may provide valuable predictions for other experimental conditions, such as investigation of cortical pain processing during pain perception in the presence of anesthetic or analgesic drugs (Zhou et al., 2018). Finally, the biophysical model can be extended as a dynamic causal model of complex cross spectral responses (Friston et al., 2012). The parameters of such a forward or generative model of observed data may be optimized using variational techniques. This will enable us to quantify both the gain or weight parameters of our model, as well as the uncertainty of these estimates. We will then be able to test hypotheses about the effects under different pain conditions.

## ACKNOWLEDGMENTS

This work was partially supported by the National Science Foundation (NSF)-CBET grant 1835000 (ZSC, JW), National Institutes of Health (NIH) R01-NS100065 (ZSC, JW), R01-MH118928 (ZSC), and a fellowship of the NIH Training Program in Computational Neuro-science (HK) supported by NIH T90/R90 DA043219 and DA043849. Preliminary version of this work was presented in Proceedings of IEEE EMBC’19, Berlin, July 23-28, 2019 (Song et al., 2019).

## AUTHOR CONTRIBUTIONS

Conceived and designed the experiments: ZSC, JW. Supervised the project: ZSC. Performed the experiments and collected the data: QZ, ZX, AS. Analyzed the data: YS, MY, HK, ZX. Contributed the software: YS, MY, AB. Wrote the paper: ZSC.

## Supplementary Materials

**Supplementary Fig. 1:**
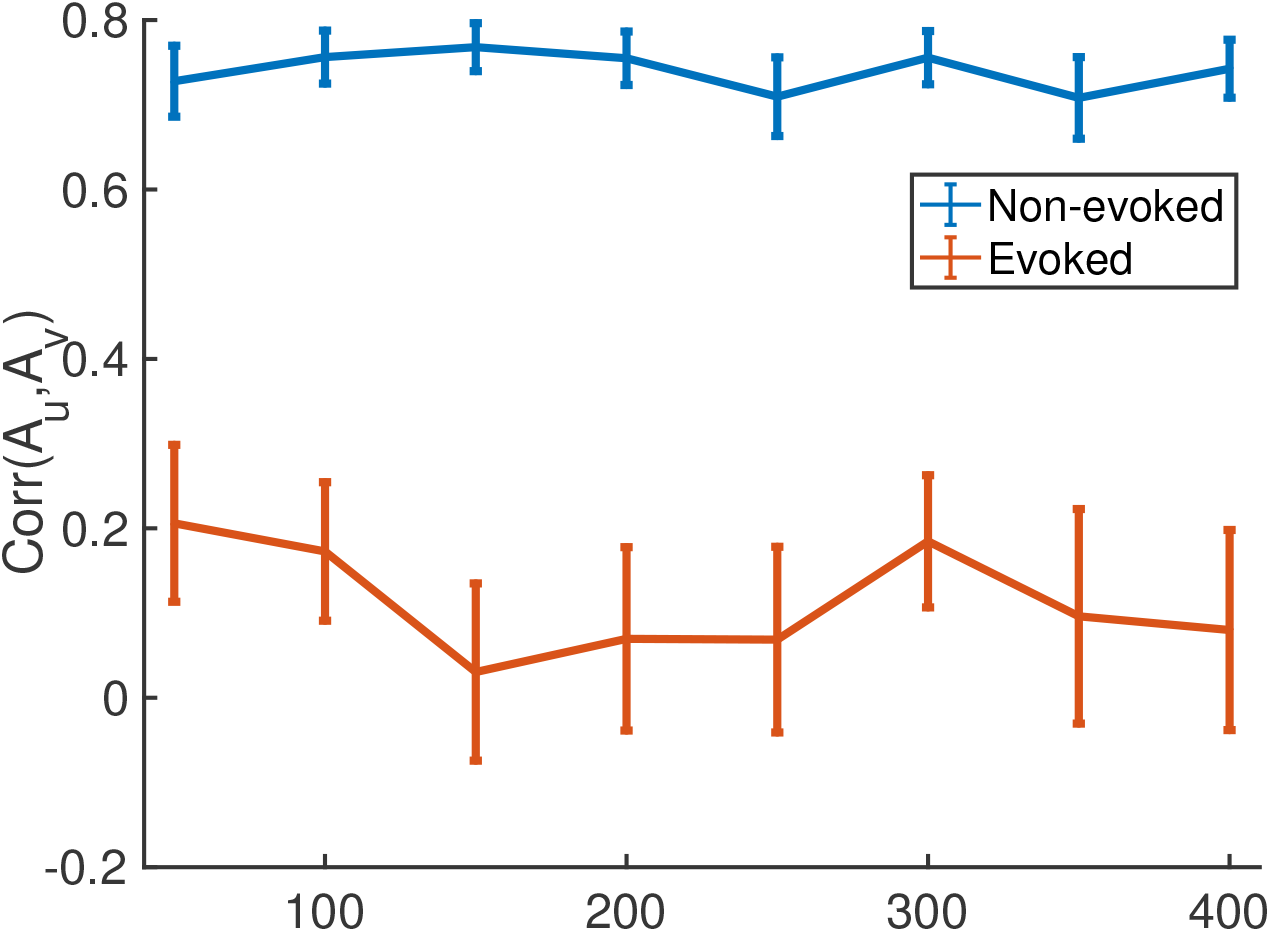
Sensitivity analysis of correlation between *A_u_* and *A_v_* with respect to the delay parameter Δ_*u*_ in the predictive coding model. The correlation statistics are relatively stable across a wide range of Δ_*u*_ in evoked pain (red) and non-evoked nociception (blue). Error bar denotes SEM (*n* = 10).

**Supplementary Fig. 2:**
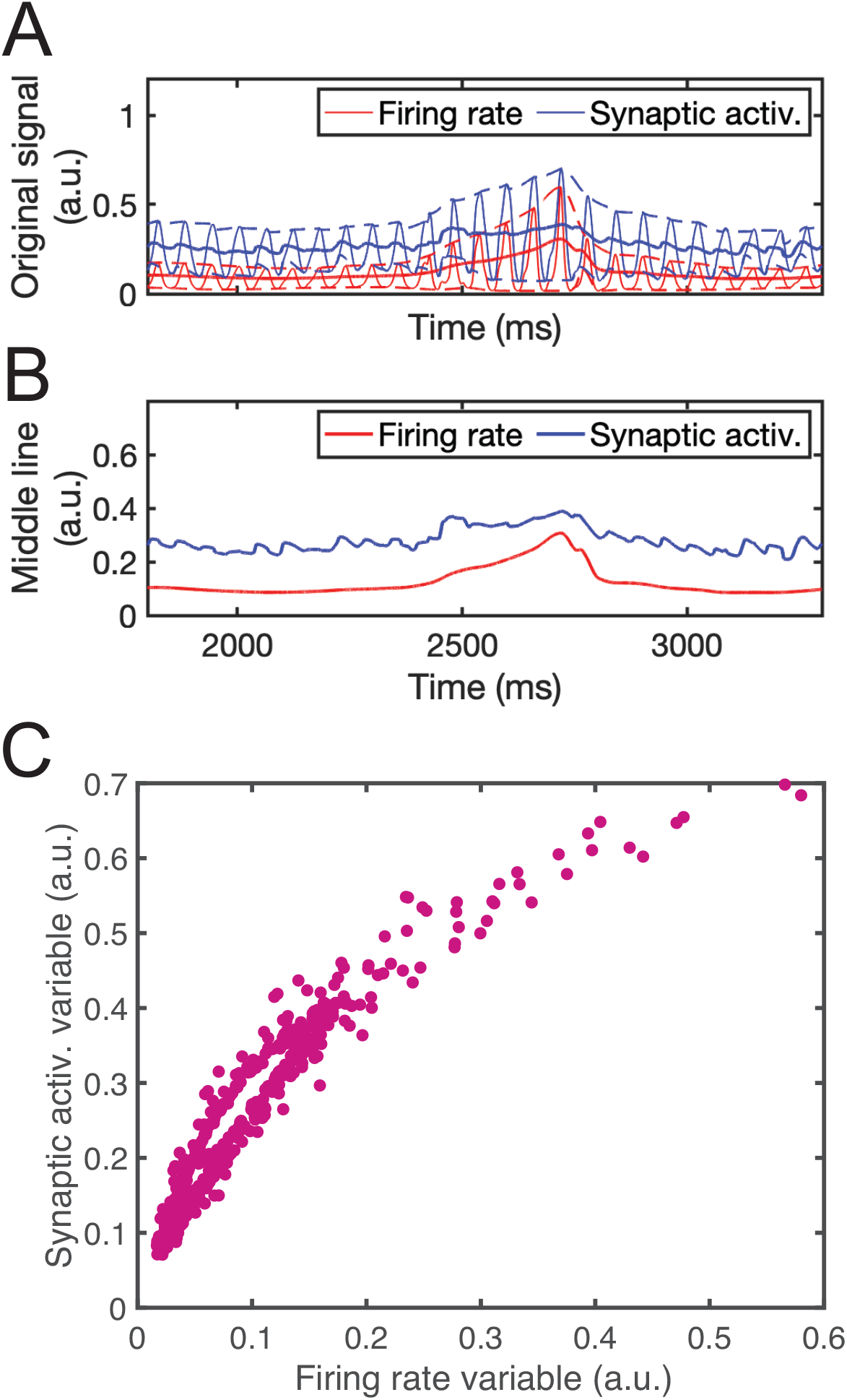
Comparison of firing rate variable *r* and synaptic activation variable *s* in evoked pain based on the mean field model. (*A*) Representative simulated traces of firing rate (red) and synaptic activation (blue) of ACC population *E*_2-1_ in one evoked pain trial. Dashed lines show the upper and lower envelope of the oscillation. (*B*) Replot the midline of the envelopes in panel A. (*C*) Scatter plot of firing rate variable and synaptic activation variable in panel A. Two variables are highly correlated (Spearman’s correlation 0.969, *p* < 10^−10^).

**Supplementary Fig. 3:**
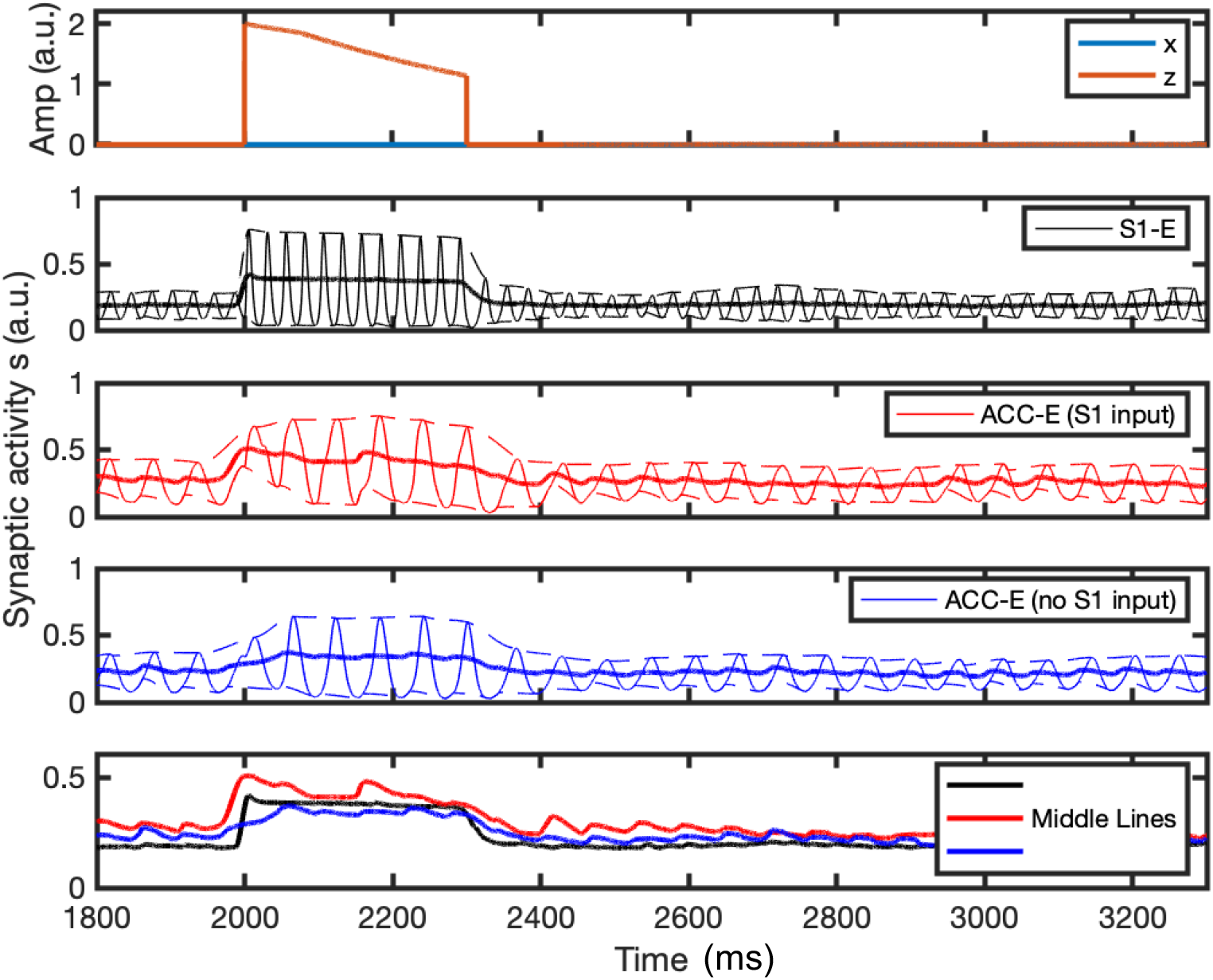
Mean-field activity (synaptic activation *s*) for three different excitatory neuronal populations in one representative non-evoked nociception simulation. Notations are the same as Fig. 6*A,B*.

**Supplementary Fig. 4:**
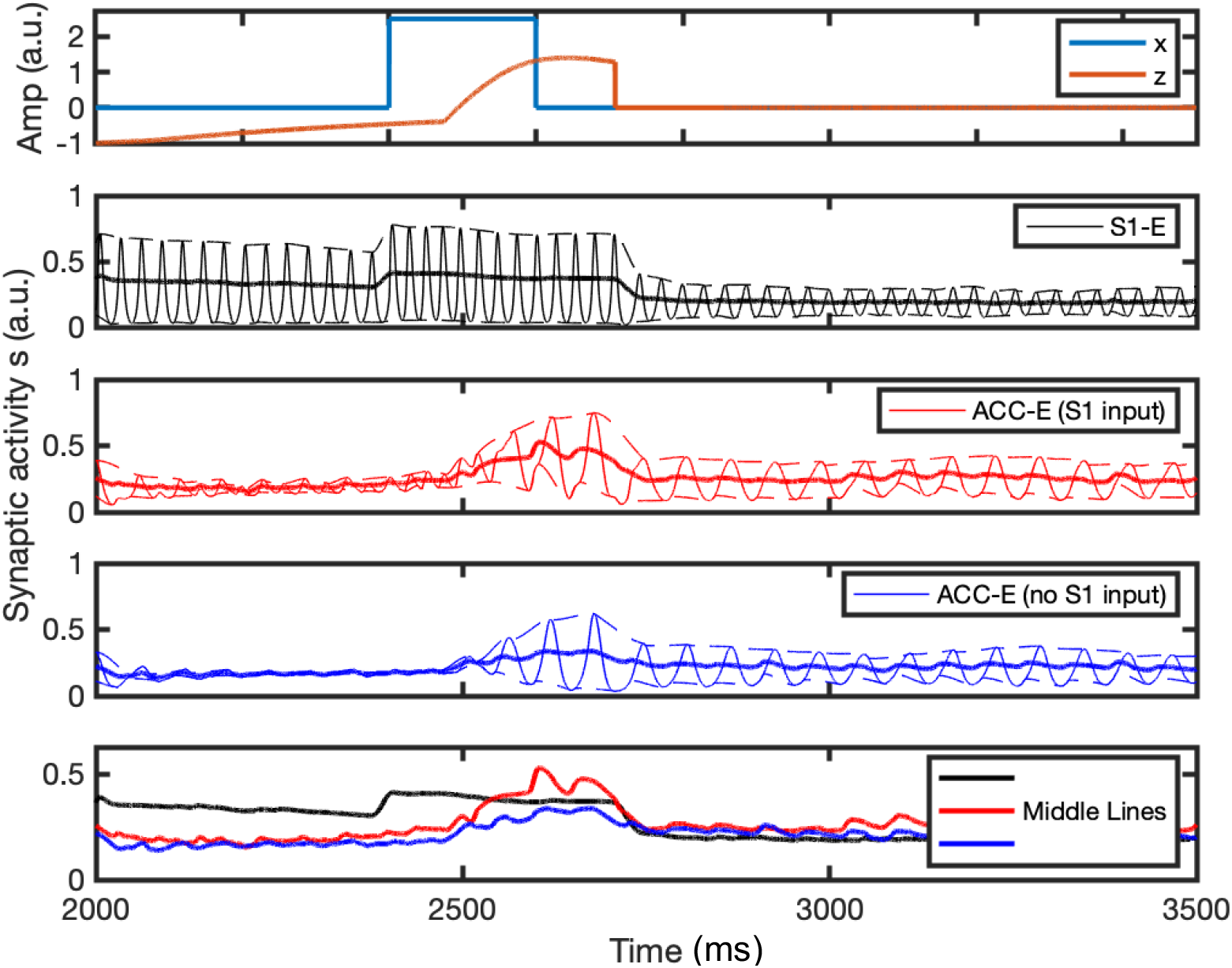
Mean-field activity (synaptic activation *s*) for three different excitatory neuronal populations in one representative placebo condition simulation. Notations are the same as Fig. 6*A,B*.

**Supplementary Fig. 5:**
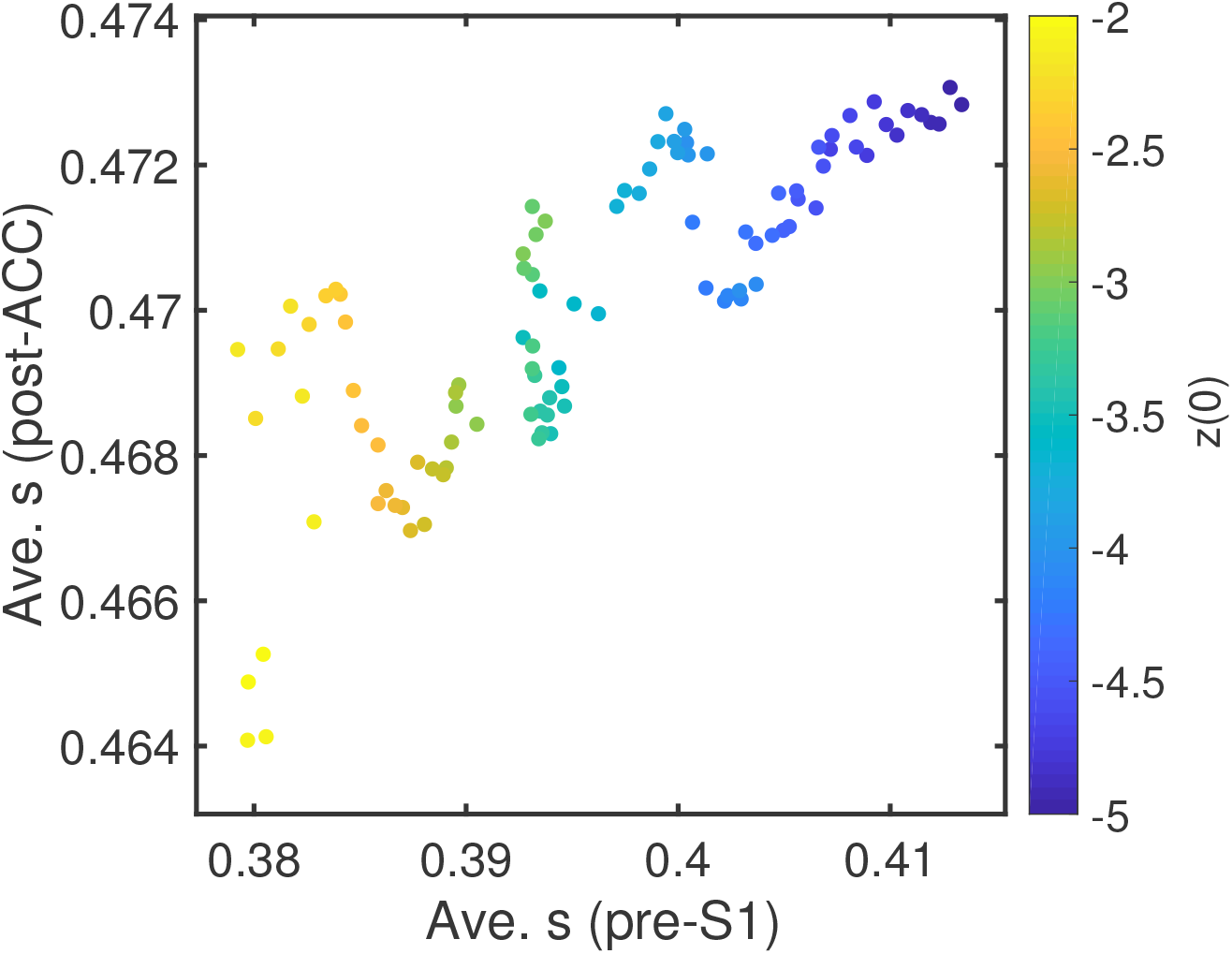
Scatter plot of the average pre-S1 synaptic activation *s* versus the average post-ACC synaptic activation *s* derived from the mean field model simulations (*n* = 100) in the placebo condition (Pearson’s correlation coefficient: 0.80, *p* = 7.7 × 10^−24^).

**Supplementary Fig. 6:**
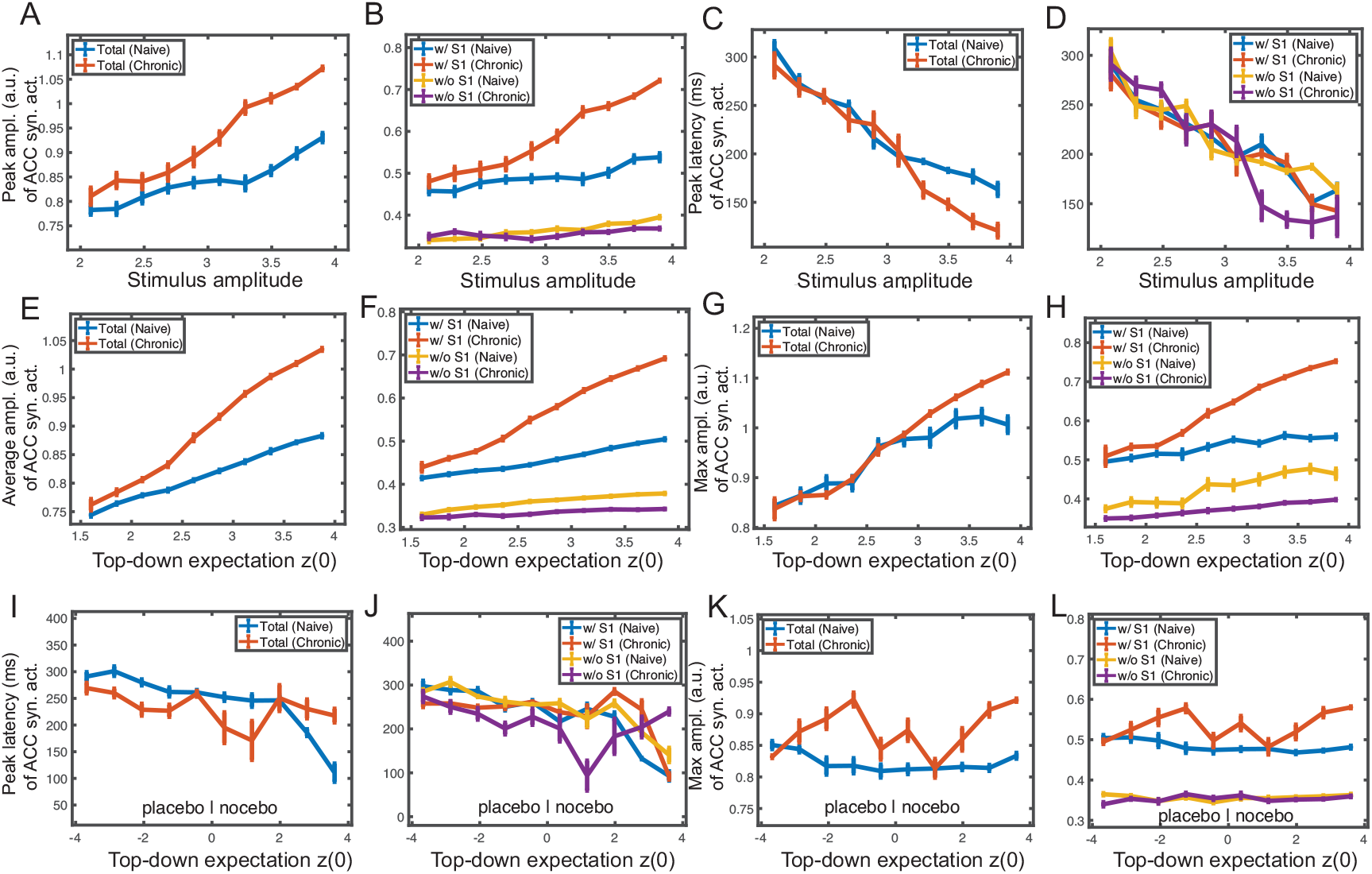
Latency and maximum peak statistics of synaptic activation in ACC populations during evoked pain. (*A-D*), non-evoked nociception (*E-H*) and placebo/nocebo (*I-L*) conditions. (*A*) Maximum of middle line of ACC synaptic activation variable *s* from total population during the duration *T_s_* between the stimulus onset to withdrawal, for varying stimulus amplitude under naive (blue) and chronic pain (red) conditions. (*B*) Similar to panel *A*, except for two ACC subpopulations. (*C*) The latency from the stimulus onset to the maximum defined in panel *A* for varying stimulus amplitude under the naive and chronic pain conditions. (*D*) Similar to panel *C*, except for two ACC subpopulations *E*_2-1_ (w/ S1 input) and *E*_2-2_ (w/o S1 input). Mean and SEM for each group are shown. 100 Monte Carlo runs were run with random initial input amplitude *x* ∈ [1.3, 5.0]. (*E*) Average of middle line of ACC synaptic activation variable *s* from the total population during the duration *T_s_* for varying top-down expectation *z*(0) under naive and chronic pain conditions. (*F*) Similar to panel *E*, except for two ACC subpopulations *E*_2-1_ and *E*_2-2_. Mean and SEM for each group are shown. 100 Monte Carlo runs were run with random initial *z*(0) ∈ [1.5, 4.0]. (*G*) Maximum of middle line of ACC synaptic activation variable *s* from total population during the duration *T_s_* between the stimulus onset to withdrawal, for varying top-down expectation *z*(0) under the naive and chronic pain conditions. (*H*) Similar to panel *G*, except for two ACC subpopulations *E*_2-1_ and *E*_2-2_. The curves in panels *G* and *H* have similar shapes as in panels *E* and *F*. (*I*) The latency from the stimulus onset to the maximum of ACC synaptic activation for varying top-down expectation *z*(0) under naive and chronic pain conditions. (*J*) Similar to panel *I*, except for two ACC subpopulations *E*_2-1_ and *E*_2-2_. Mean and SEM for each group are shown. 100 Monte Carlo runs were run with random initial *z*(0) ∈ [−4.0, 4.0]. (*K*) Maximum of middle line of ACC synaptic activation variable *s* from the total population during the duration *T_s_* between the stimulus onset to withdrawal, for varying top-down expectation *z*(0) under naive and chronic pain conditions. (*L*) Similar to panel *K*, except for two ACC subpopulations *E*_2-1_ and *E*_2-2_.

**Supplementary Fig. 7:**
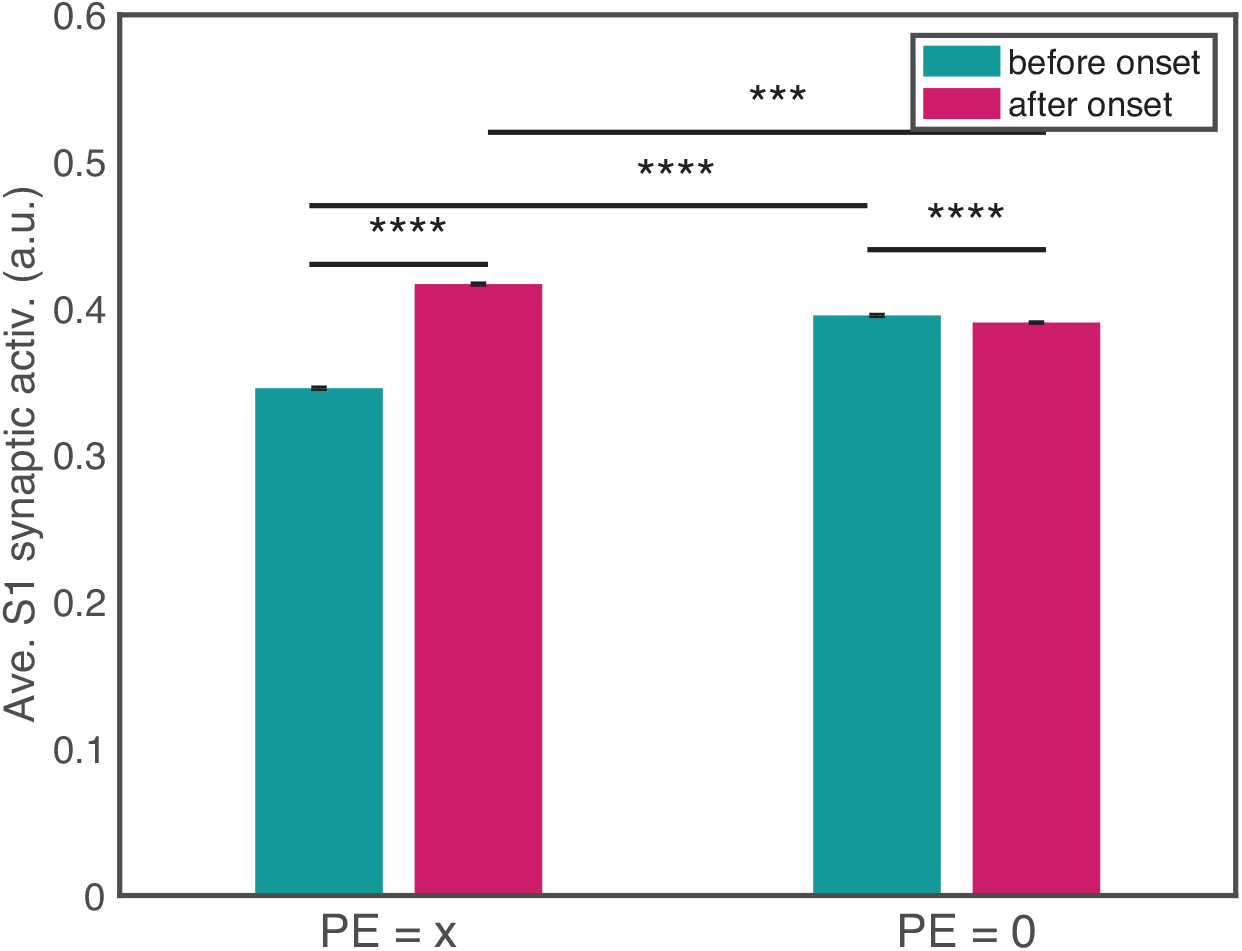
Comparison of average S1 synaptic activation at different periods (before vs. after onset) and PE values: PE= *x* (or *z* = 0) and PE= 0 (or *z* = *x*), with feedback from the ACC to S1. A total of 10 Monte Carlo trials were run with random stimulus input amplitude *x* ∈ [1.8, 2.2]. Mean and SEM were presented for each group. There was a significant difference in the average S1 synaptic activation variable between before and after the stimulus onset in both conditions. All *p*-values for pairs marked in the graph are less than 0.0001, expect for the *p* = 0.0008 between PE= *x* and PE= 0 after the onset (two pink bars). This indicates that the decrease in S1 firing intensity after the stimulus onset was slightly less significant with the presence of feedback. The pre-stimulus firing was averaged from the expectation *z* onset (from 0 if no expectation) to the stimulus *x* onset; the post-stimulus firing was averaged from the stimulus onset to withdrawal. Compared to Fig. 9*C*, the gap between before and after the stimulus onset was smaller here.

